# A highly-parallel microchannel platform reveals the transcriptomic and epigenetic signature of confined migration

**DOI:** 10.64898/2026.11.15.766750

**Authors:** Yixuan Li, Lee Jia Wen Nicole, Xing Chen, Chang Ye Ji, Vaishnavi Rangaraj, Malhar S. Chitnis, Zhou Fang, Sriram Muthukumar, Shinny Sunny, Kabigting Jessica Evangeline Tan, He Yuxuan, Fu Zhehui, Jennifer Marlena, Apoorva Shivankar, Chii Jou Chan, Zhu Jin, Yusuke Toyama, Andrew W. Holle

## Abstract

Confining tissue microenvironments are ubiquitous in vivo, yet existing in vitro platforms are limited in geometric control, throughput, or compatibility with post-migration cell recovery. We developed a highly parallel microchannel membrane containing thousands of tunable microchannels in a format compatible with standard cell culture, imaging, cell retrieval, and direct lysis. The platform enables control over channel cross-sectional area, length, and geometry across multiple cell types. Using this system, we recovered sufficient post-confined cells for bulk RNA-seq and ATAC-seq, revealing confinement-dependent transcriptomic and chromatin accessibility signatures. Increasing confinement was associated with predominantly reduced chromatin accessibility, while comparison of two cancer cell lines identified shared differentially expressed genes and convergent enrichment of cancer-associated, inflammatory, and EMT-related pathways. This scalable platform enables omics-scale analysis of cells after migration through defined physical microenvironments.

## Introduction

Confining microenvironments are prevalent *in vivo*, particularly within interstitial spaces where cells migrate through densely packed tissues and matrix-rich regions^1–3^. These physically confining regions are encountered during key biological processes such as development, immune cell homing, wound healing, and cancer invasion^4–7^. Confinement imposes spatial constraints that require cells to undergo dramatic morphological adaptations^8^, such as nuclear deformation^9–12^ and cytoskeletal reorganization^8,13,14^, which in turn influence gene expression and long-term cell fate^15^. Recent studies have shown that confined migration can drive stem cell differentiation and alter migratory persistence, chromatin architecture, and transcriptional programs^16,17^. These findings highlight the need for experimental systems that can more effectively model and interrogate the effects of confinement on cells.

To study the effects of physical confinement *in vitro*, a range of engineered systems have been developed, including sandwich confinement assays, microchannel devices, and transwell inserts^18,19^. These platforms have significantly advanced our understanding of how cells sense and respond to confined spaces^20,21^. By recapitulating key aspects of the *in vivo* microenvironment, they offer control over dimensionality, extracellular matrix composition, and flow, while remaining compatible with co-culture setups and downstream analyses such as immunostaining^22–25^. This makes it possible to explore how confinement shapes cell polarity, migration modes, nuclear mechanics, and signaling pathways in a controlled setting^8,26–28^.

Despite these strengths, current *in vitro* confinement platforms face several practical and technical limitations that constrain their broader utility. Microchannel-based systems, for example, are typically closed microfluidic devices that are not designed for efficient post-confinement cell retrieval. The few platforms that are designed to enable cell retrieval require increasingly specialized geometries and fabrication processes, necessitating complex fluidic circuits or harsh retrieval protocols for on-chip lysis^29–31^. Moreover, the throughput of these systems is generally low, making it challenging to collect sufficient cell numbers for bulk RNA seq, ATAC seq, or protein-based assays^15,32,33^. While sandwich confinement assays have been widely applied to investigate cellular responses to mechanical compression and are compatible with bulk analysis, confinement in these systems is externally imposed on cells, whereas confined migration systems require cells actively entering physiologically tight spaces. In addition, prolonged vertical confinement has been reported to induce programmed cell death^34^. These two experimental paradigms therefore represent distinct mechanical contexts and may elicit different cellular responses.

While traditional transwell inserts offer better scalability, they present a different set of limitations. Their thin membranes do not permit long-range cell migration, preventing the study of dynamic processes over extended confinement distances that mimic interstitial spaces. Additionally, the architecture of the pores lacks precise geometric control, limiting the ability to model defined confinement conditions with spatial accuracy^35^. These technical constraints collectively limit the biological questions that can be addressed using current confinement tools.

Here, we have developed a high-throughput confinement system that enables both precise geometric control of confined environments and efficient recovery of cells or lysates for downstream analysis. Using this system, we observed distinct confinement-dependent cell morphologies, including drastic cytoskeletal adaptation and nuclear deformation in response to severe confinement and unique cell-ECM interactions at the 2.5D edges of wide microchannels. Most importantly, the scalable nature of this platform enabled us to collect large numbers of post-confinement cells, allowing for transcriptomic profiling of the response to confinement. Our system provides a new window into the transcriptomic and epigenomic signature of physiological confinement and can facilitate new clinical approaches to disease that aim to leverage both the proper and aberrant cellular response to confinement.

## Results

### Confinement membrane fabrication with high aspect ratio microstructures

To overcome the inability of traditional photolithography and replica molding to produce arrays of high aspect ratio structures, a new fabrication method was developed to produce PDMS membranes with tunable arrays of microchannels spanning the membrane plane (**Fig. 1a**). Briefly, standard photolithography was used to produce master molds containing rows of individual microchannel structures. Uncured PDMS was then spin-coated onto these wafers and removed as thin films with the aid of a thicker handling layer and plasma-activated PDMS-PDMS attachment. This process was repeated seven additional times, yielding a PDMS block containing eight layers of microchannels. Individual blocks were then vertically stacked to further increase the quantity of microchannels in the cross-sectional surface. Finally, these assembled PDMS blocks were sliced with a vibratome into 150 µm sheets, yielding large numbers of membranes containing thousands of transmembrane microchannels, enabling thousands of cells to migrate through the channels in parallel (**Extended Data Fig. 1a-c**). In this approach, the photolithographically patterned structures define the channel cross-section, while the final channel length is determined by the thickness of the vibratome-cut membrane, allowing independent control over both parameters (**Extended Data Fig. 2**)

**Fig. 1.**
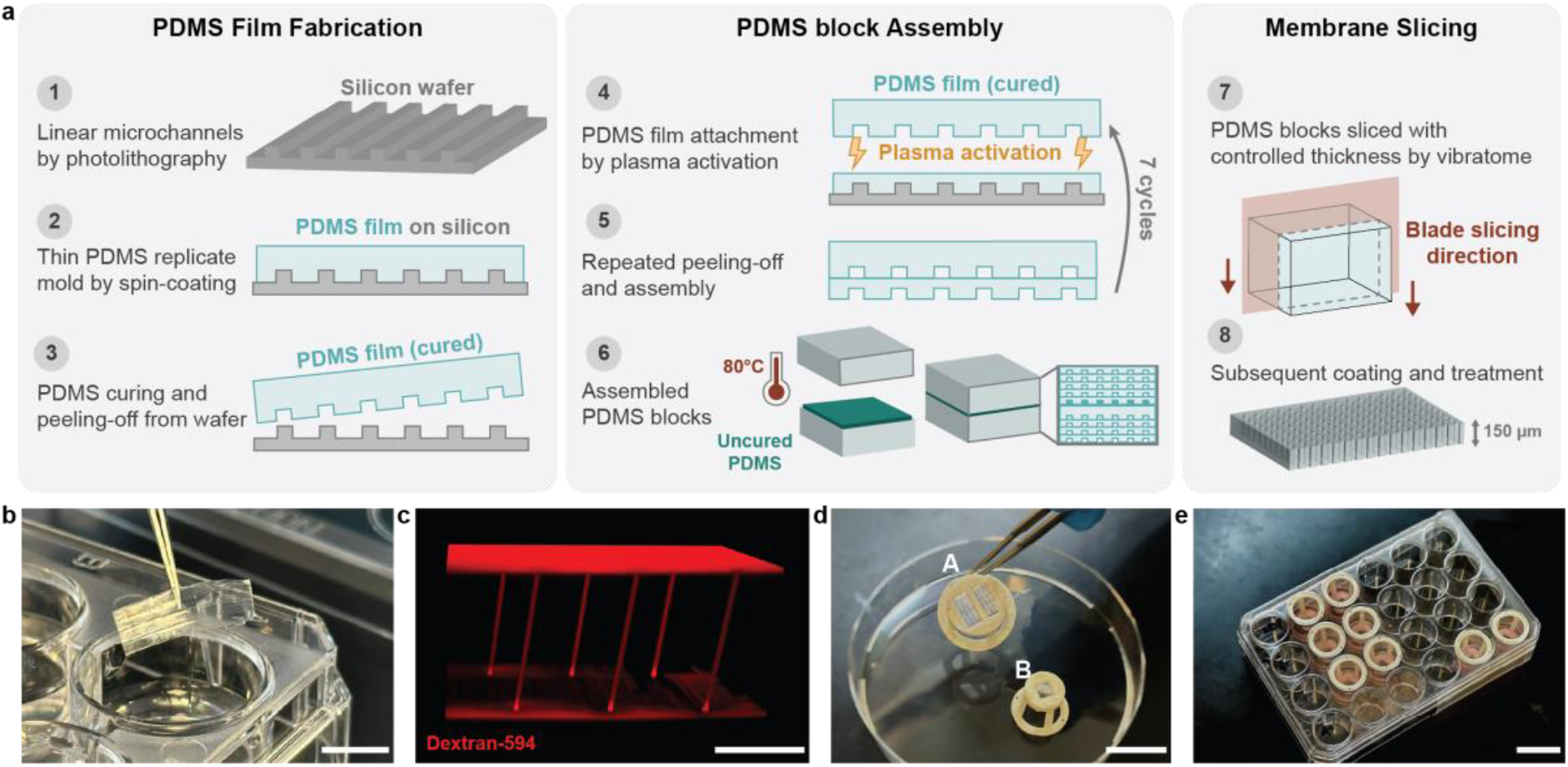
| Fabrication process and functional principles of the microchannel membranes. **a**, Fabrication workflow: PDMS film fabrication (left), plasma-activated assembly of layered PDMS block (middle), and final porous microchannel membrane production (right). **b**, Microchannel membrane post-slicing. Scale bar: 1 cm. **c**, 3D reconstruction of fluorescent dextran-labelled microchannels (30 µm² × 150 µm) within a membrane. Scale bar: 100 µm. **d**, Custom resin inserts designed to secure microchannel membranes for 12 well (A) and 24 well (B) plates. Scale bar: 2 cm. **e**, Cell culture setup with microchannel membrane-integrated inserts in a standard 24 well plate. Scale bar: 2 cm.

The resulting microchannel membranes were optically clear (**Fig. 1b**), biocompatible, and suitable for surface modification such as protein coating or passivation. Commercial transwell assays, which claim to provide similar levels of axial confinement, were found to have uneven spatial distribution of channels, resulting in large, merged pores of irregular shapes that cannot impose consistent cell and nuclear deformation (**Extended Data Fig. 3d-h**). Additionally, transwell assay membranes were found to be less than 10 µm thick (likely due to limitations in fabrication method), which is not sufficient to fully contain either the cell body or even the nucleus on its own (**Extended Data Fig. 3i and j**). In comparison, our method produces microchannel structures with extremely high aspect ratios and precise spatial patterning (**Fig. 1c, Extended Data Fig. 1a**). For the 150 µm-long channels used here, this corresponds to length-to-height aspect ratios of approximately 30:1, 15:1, and 7.5:1 for the high, low, and wide channels, respectively. By slicing the assembled PDMS blocks orthogonally, we have precise control over the length of the microchannels within the array, allowing for the production of membranes with continuous confining microchannels of lengths ranging from approximately 50 µm up to 2 cm for specialized applications. Moreover, because photolithography-based patterning allows for precise control over microscale features, we can go beyond the fabrication of standard linear microchannels and produce a range of channel geometries within the PDMS membrane. In addition to the 30, 100, and 400 µm² channels used throughout this study, we fabricated channels with cross-sectional areas as small as 9 µm², demonstrating the ability of the platform to generate more extreme confinement conditions (**Extended Data Fig. 4h**). This approach can also be used to generate irregular and interconnected structures that mimic the branching morphology found in tissue microenvironments, including capillary networks, as well as periodic confinement channels containing repeated constrictions along the length of the channel (**Extended Data Fig. 4a-e**). The channel length can also be tuned independently during the vibratome slicing step. While 150 µm-long channels were used for the majority of subsequent experiments, we also fabricated membranes containing 50 µm and 200 µm-long channels, demonstrating that final channel length is not fixed by the photolithographic mold but can instead be adjusted during membrane sectioning (**Extended Data Fig. 4f and g**).

To enable confocal imaging with subcellular resolution and seamless integration with standard workflows, customized resin inserts that fit into conventional cell culture plates were designed and fabricated (**Fig. 1d**). PDMS membranes were affixed to these inserts using either uncured PDMS or silicone glue. Uncured PDMS provides a strong, permanent bond suitable for long-term cultures or mechanical stability, while the silicone glue allows for easy removal after culture, facilitating downstream immunofluorescence experiments. The resulting setup supports routine procedures such as media exchange, fixation, and co-culture, all within a standard culture system (**Fig. 1e**).

### Controlled production of microchannels across a wide range of sizes

To recreate different degrees of physiological confinement *in vitro*, microchannel membranes with cross-sectional areas of 400 µm^2^, 100 µm^2^, and 30 µm^2^ were fabricated and coated with collagen (**Fig. 2a**). The 3D orientation and length of the channels were measured to verify the accuracy and the reproducibility of the fabrication process (**Fig. 2b** **and Extended Data Fig. 3a-c**), with channel length approximately 150 µm and vertical tilt consistently below 7° (**Fig. 2c and d**). While this slight tilt does not cause a notable change in overall channel length, it does lead to minor optical effects that reduce fluorescence intensity toward the middle of the channels, as also observed with fluorescent dextran (**Extended Fig. 4i-k**). Quantification of width, height, and cross-sectional area showed close agreement with the intended designs for all three channel types (**Fig. 2e-g**).

**Fig. 2.**
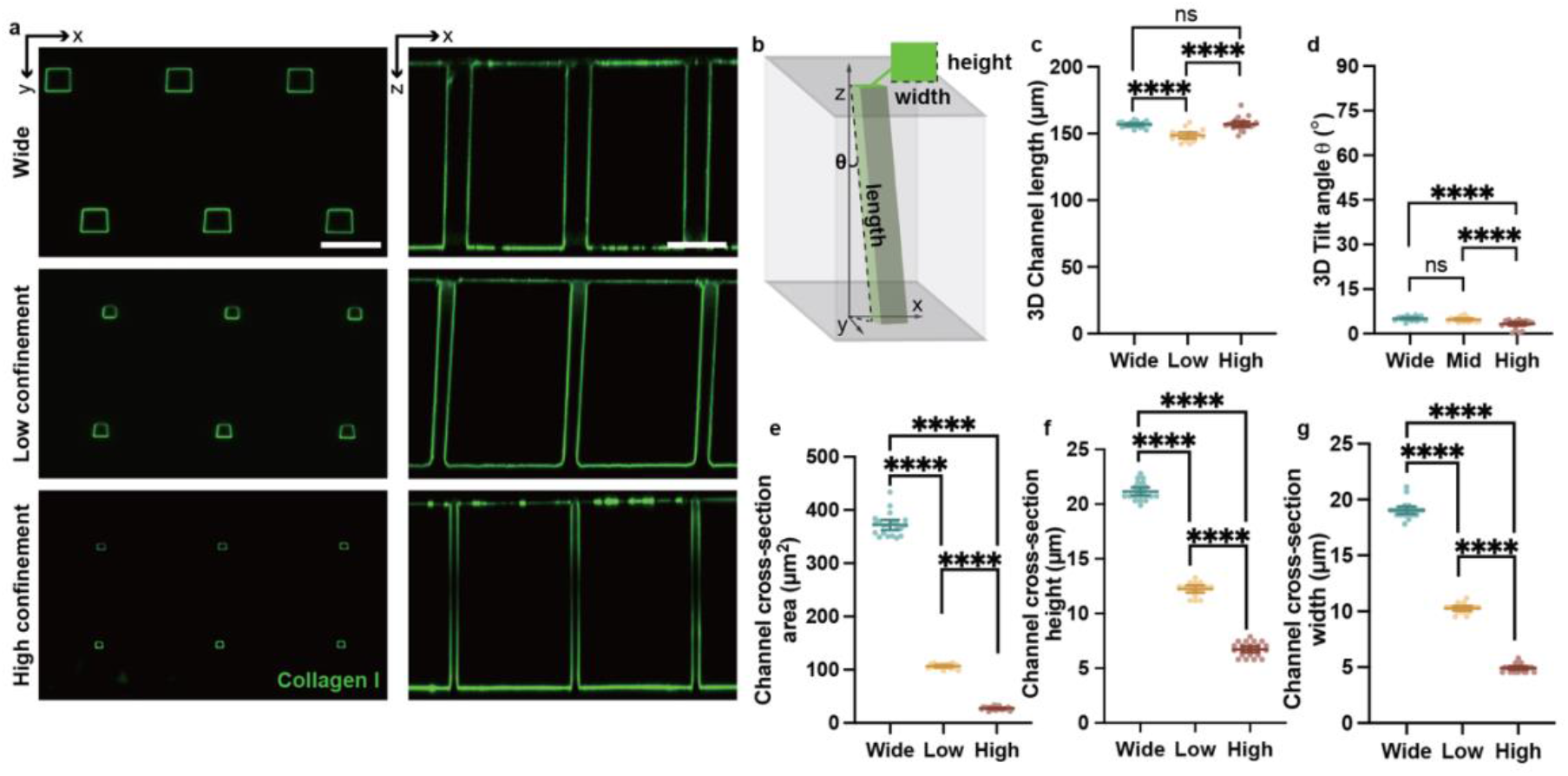
| Structural characterization of sliced membranes across confinement sizes. **a**, Representative fluorescence images of wide, low-confinement, and high-confinement microchannels in x-y (left) and x-z (right) dimensions. Scale bar: 50 µm. **b**, A diagram showing the measured 3D features of the trans-membrane microchannels. **c–d**, Quantification of 3D microchannel length (c) and tilt angle (d). **e–g**, Quantification of microchannel cross-sectional area (e), height (f), and width (g). Statistical comparisons: one-way ANOVA with Holm-Šídák post-hoc test. Significance: ns (not significant), *P ≤ 0.05, **P ≤ 0.01, ***P ≤ 0.001, ****P ≤ 0.0001. Error bars: mean ± 95% C.I. (Wide: n=21, Low: n=17, High: n=22).

### Confinement-induced morphological plasticity in migratory cancer cells

HT1080 cells, despite being originally derived from fibrosarcoma tissue, are considered to have a relatively stable genome^36^. Due to their migratory capacity, they serve as a widely utilized model system for investigating key processes in cancer metastasis, including matrix remodeling, adhesion, and cell motility^37–41^. On control 2D PDMS surfaces, HT1080 cells exhibited well-defined actin stress fibers, broad lamellipodia, and multiple regions of thin filopodia (**Fig. 3a**). Within wide microchannels that do not impose confinement, cells exhibited thin protrusive structures that interacted with multiple walls of the microchannels (**Fig. 3a-b**).

**Fig. 3.**
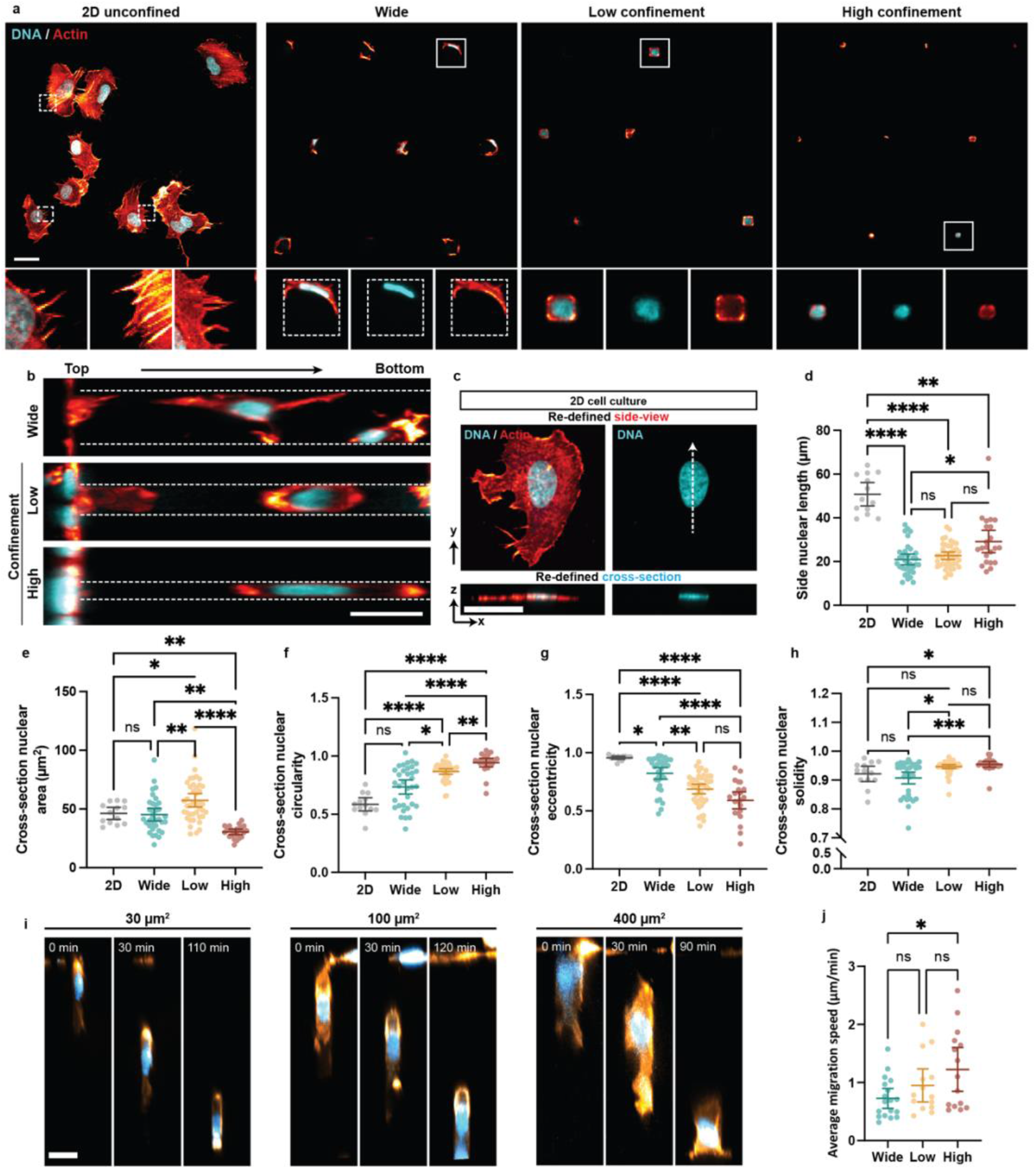
| Cytoskeletal architecture and nuclear remodeling during confined migration through the microchannel membrane. **a**, Cytoskeletal structure and nuclear morphology of HT1080 cells cultured on 2D PDMS membranes and migrating through wide, low-confinement, and high-confinement channels in the microchannel membranes. Zoomed-in regions highlight individual cells at microchannel cross-sections (dashed lines: channel boundaries). **b**, Side-view images of transmigrating cells, showing cytoskeletal organization and nuclear morphology (dashed lines: microchannel boundaries). **c**, The definition of cell orientation of 2D-cultured cells for morphology measurements. **d–h**, Nuclear morphology quantified from side-view (d) and cross-sectional (e–h) images: nuclear major axis length from side-views (d); nuclear area (e), circularity (f), eccentricity(g), and solidity (h) from cross-sections (Wide: n=34, Low: n=42, High: n=22). Statistical comparisons: one-way ANOVA with Holm-Šídák post-hoc test (e) and Kruskal-Wallis test with Dunn’s post-hoc test (d, f–h). Significance: ns (not significant), *P ≤ 0.05, **P ≤ 0.01, ***P ≤ 0.001, ****P ≤ 0.0001. Error bars: mean ± 95% C.I. Scale bars: 30 µm. **i**, Representative timelapse images of HT1080 cells migrating through wide, low, and high confinement microchannels. Scale bars: 20 µm. **j**, Quantification of cell migration speed through wide, low, and high confinement channels. (Wide: n=18, Low: n=14, High: n=15). Statistical comparisons: Kruskal-Wallis test with Dunn’s post-hoc test. Significance: ns (not significant), *P ≤ 0.05, **P ≤ 0.01, ***P ≤ 0.001, ****P ≤ 0.0001. Error bars: mean ± 95% C.I.

To better understand 3D nuclear deformation in microchannels, we quantified nuclear morphology, both perpendicular and parallel to the channel axis. For control 2D substrates, the perpendicular projection was defined as the z-plane orthogonal to the substrate, and the parallel projection was defined as the x-y projection, mimicking the spatial orientation of confined cells (**Fig. 3c**). Interestingly, nuclei within wide control microchannels were significantly shorter, more flattened, and slightly concave compared with 2D controls (**Fig. 3d-h**). These differences indicate that exposure to the three-dimensional channel geometry is associated with altered nuclear morphology even in the absence of direct physical confinement.

In both low confinement (100 µm²) and high confinement (30 µm²) microchannels, HT1080 cells and their nuclei were heavily deformed, to a degree that they completely occupied the available cross-sectional area (**Fig. 3a**). In low confinement, prominent actin signal was observed along the cell periphery (**Fig. 3a**) in close apposition to the channel walls, consistent with previously observed changes in cortical actin organization under confinement. Quantification of phalloidin intensity along the migration axis revealed actin enrichment toward both the leading and trailing regions of cells in the low confinement (**Extended Data Fig. 5c**). Under high confinement, F-actin organization became more strongly polarized, with pronounced enrichment toward the trailing edge of cells in the high confinement, consistent with previous reports of rear-biased cortical actin organization and contractility during confined and amoeboid migration^42–44^ (**Extended Data Fig. 5d**). Nuclei in the low confinement channels were elongated but retained a relatively smooth contour, whereas nuclear deformation increased under high confinement, with cells exhibiting highly elongated and narrow nuclei aligned with the channel axis (**Fig. 3b, d-h**). In general, as confinement increased, nuclei became more elongated along the channel axis and more round in cross-section (**Fig. 3d, f-h**), mirroring observations made in standard microchannel assays^3,10,13,16,27^. To directly examine migration dynamics within the microchannels, we performed live-cell imaging of HT1080 cells traversing the wide, low-, and high-confinement channels. Migration speed varied with confinement, with cells migrating fastest through the high-confinement channels and significantly faster than in wide channels, mirroring previous publications using standard microchannels^10,13,16,42,45,46^ (**Fig. 3i and j**). The difference in average traversal time between cells migrating through the high- and wide-channel conditions was approximately 90 minutes. Because cells were collected after a 48 hour migration period and entered and exited the channels asynchronously, the sequencing measurements reflect population-averaged post-migration states with variable time elapsed since channel exit.

Beyond channel entry and subsequent confined migration, exit from microchannels represents a key transitional state where cells re-adapt to a less confining microenvironment. To understand how cells transition out of extreme confinement, we imaged post-migratory HT1080 cells as they exited the 30 µm² channels. These cells exhibited distinct intermediate morphologies, including partially emerged states where the nucleus remained constricted at the channel interface (**Extended Data Fig. 5a)**. Notably, during this transition, cells displayed a pronounced nuclear bending, with one end remaining in the channel aligned along the channel axis and other exiting onto the 2D surface. Immunofluorescence revealed a pronounced accumulation of phosphorylated myosin light chain (p-MLC) and paxillin at the leading edge (**Extended Data Fig. 5b**), reflecting localized traction force generation and adhesion remodeling during emergence. These observations suggest that rather than passively exiting confinement, cells must exert considerable contractile forces and rearrange their cytoskeleton in order to adapt to the new microenvironment.

### A versatile system for probing mechanical adaptation across diverse cell types

A large body of literature on confinement focuses on cancer cells due to their mesenchymal-like behavior^13^, high motility^10,47^, and close connection to tumor progression and metastasis^2,12,48–50^. Beyond cancer, confined migration in physiological tissue microenvironments has been shown to induce significant changes in a wide range of cells and therefore approaches compatible with high throughput analysis of confinement in these diverse cell types are of great importance to the field. Despite large differences in cell spread area, nuclear area, and nuclear morphology (**Fig. 4a-e, Extended Data Fig. 6**), we found that both human mesenchymal stem cells (hMSCs) and macrophages were capable of entering and migrating through both low confinement and high confinement microchannels (**Fig. 4f and g**). As these cross-sectional areas are representative of the dimensions of tissue tracks and interstitial spaces^51^ encountered during processes such as stem cell homing and immune surveillance, this underscores not only the relevance of our system to diverse biological contexts, but also its ability to accommodate a broad range of cell types with distinct physical properties. Interestingly, while HT1080 cells and macrophages showed similar patterns in migration and nuclear morphology, hMSCs exhibited a distinct nuclear morphology that was more compressed and slender than its counterparts, even leaving parts of the cross-sectional channel area unfilled in some instances (**Fig 4f**). This cell type-specific response to confinement highlights the need for tools to assess these adaptations in a systematic fashion.

**Fig. 4.**
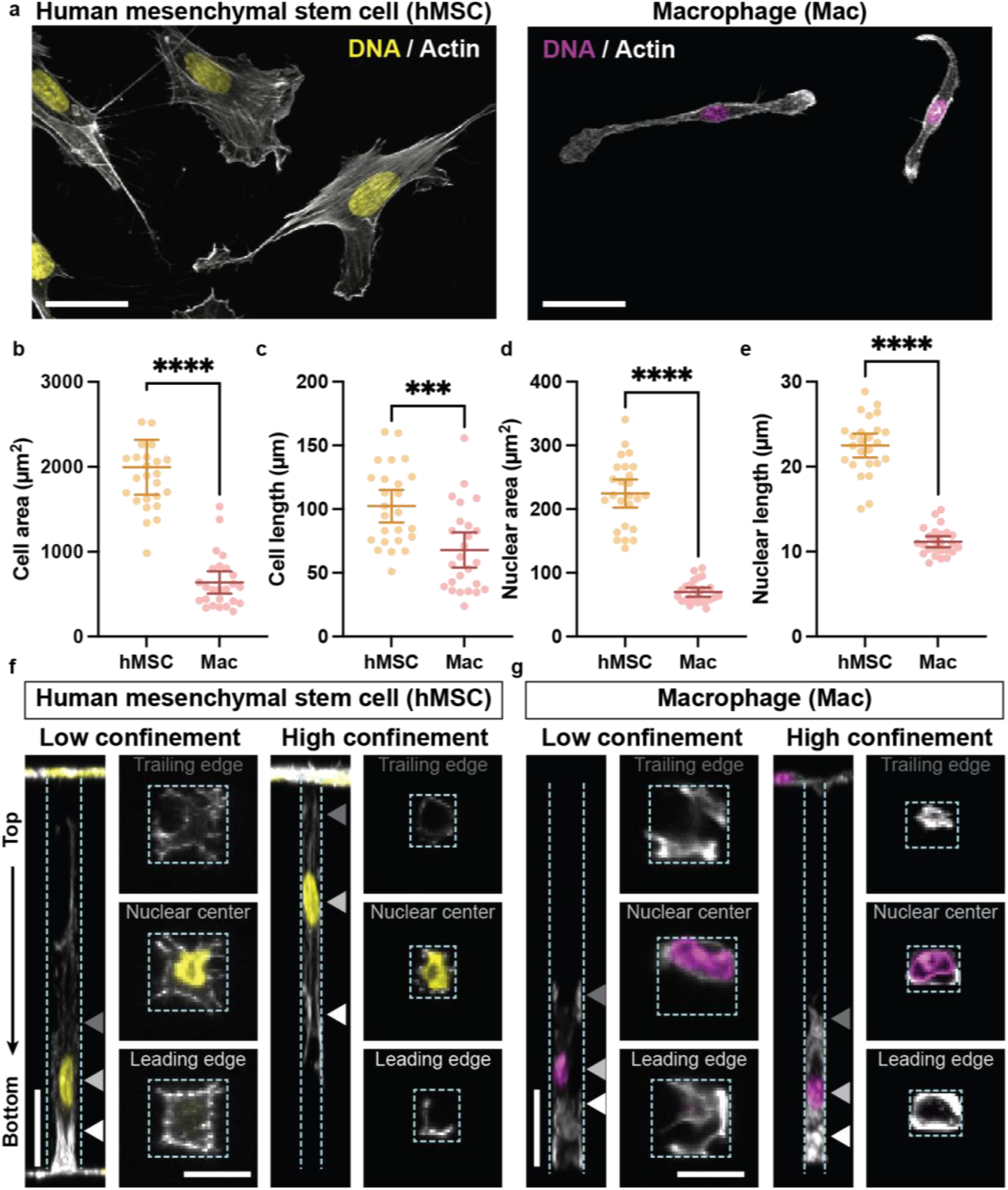
| Broad compatibility of the microchannel membrane systems across diverse cell types. **a**, Human mesenchymal stem cells (left) and primary mouse macrophages (right) morphology on PDMS. Scale bars: 30 µm). **b-e**, Quantification of cell area (b), cell length (c), nuclear area (d) and nuclear length (e) of hMSCs and macrophages on PDMS (hMSC: n=25, Mac: n=25). One outlier data point was outside the axis of b. Statistical comparisons: Mann-Whitney test (b, c) and Welch’s t test (d, e). Significance: ***P ≤ 0.001, ****P ≤ 0.0001. Error bars: mean ± 95% C.I. **f** and **g**, Cell and nuclear morphologies of hMSCs and macrophages under low and high confinement. Scale bars: 30 µm. Zoomed-in regions highlight the trailing edge, nuclear center and leading edge of individual cells at microchannel cross-sections. Scale bars: 10 µm. White arrows: approximate cross-section positions. Dashed lines: channel boundaries.

### Unique transcriptomic features resulting from physical confinement

Previous studies have demonstrated that physical forces can alter cell behavior and morphology^12,17^. While different mechanistic pathways have been identified to explain the translation of physical forces into molecular signals, little is known about global transcriptional and chromatin accessibility changes triggered by physical confinement. This is primarily due to technical challenges in collecting sufficient material from post-confinement cells using conventional microfluidic platforms^52^. In this system, culturing cells on the upper surface of a microchannel membrane with a surface area of one square centimeter for 48 hours yields a large, confinement-dependent number of post-confinement cells compatible with trypsinization or lysis. Consistent with previous reports of larger numbers of cells migrating through low levels of microchannel confinement compared to highly confining microchannels, we were able to collect ∼100,000 cells from a single low confinement (100 µm^2^) membrane and ∼5,000 cells from a single high confinement (30 µm^2^) membrane (**Extended Fig. 7e**). In wide control membranes that do not impose any confinement (400 µm^2^), the comparatively lower ∼25,000 cells reflected the fact that as these channels are wider than the diameter of a suspended cell, many seeded cells fell through the channels, a gravity-driven process that does not occur in low confinement or high confinement membranes. As the assay does not distinguish cells that attempted to enter a channel but ultimately failed from those that did not initiate entry, these measurements cannot be used to determine the fraction of cells that initiate migration or the specific factors driving channel entry.

We performed bulk RNA-seq on post-migration HT1080 cells that had navigated wide control channels (400 µm²), low confinement channels (100 µm²), and high confinement channels (30 µm²), as well as unconfined 2D control cells. Importantly, cells recovered from the wide, low, and high confinement conditions had all actively migrated through the membrane, allowing comparisons between channel conditions to distinguish molecular responses associated with increasing confinement from those associated with migration itself. Transcriptome analysis revealed that high confinement produced the largest number of differentially expressed genes (DEGs; 112 genes), indicating a strong transcriptional response to extreme spatial confinement (**Fig. 5a**). In contrast, low confinement condition elicited fewer DEGs (37 genes), suggesting a milder molecular shift that positions it as an intermediate state between 2D and high confinement (**Fig. 5a**). Interestingly, migration through wide control channels also resulted in a substantial number of DEGs (80 genes) compared to 2D control substrates (**Fig. 5a**). In view of the distinct cell morphology observed in this 3D microenvironment (**Fig. 3a**), we concluded that the wide control channels elicit a separate 3D migration mode in these cells, as opposed to being functionally identical to 2D substrates.

**Fig. 5.**
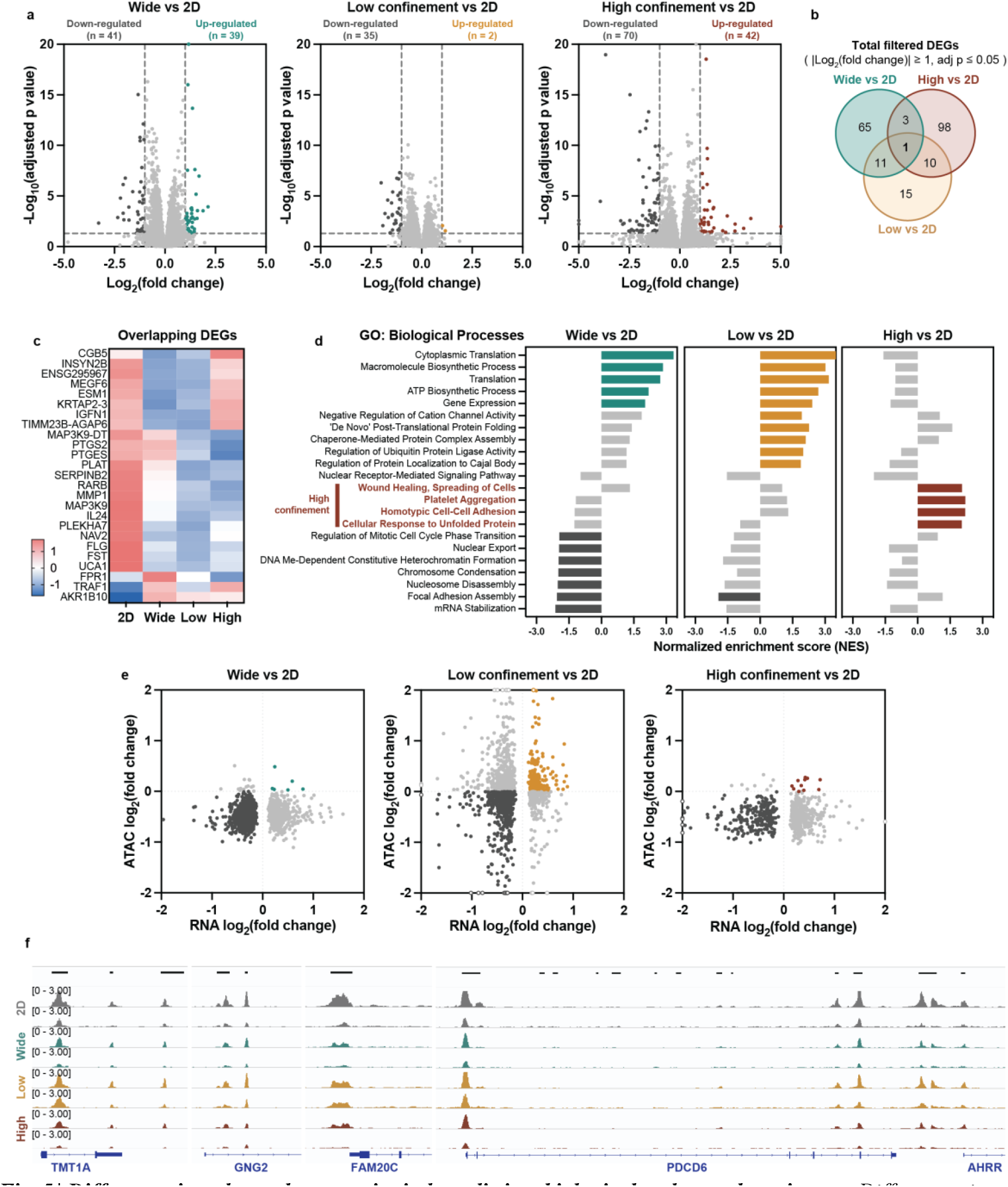
| Different microchannel geometries induce distinct biological pathway alterations. **a**, Differences in gene expression between post-confinement cells and 2D cultured cells. **b**, Venn diagram of DEGs post different degrees of confinement. Numbers indicate the number of DEGs present in each group. **c**, Heatmap showing the z-scores DEG expression shared between different confinement groups. **d**, GO (Biological Processes) gene set enrichment analysis. High-confinement specific terms were highlighted in red, and light gray color marks the terms with False Discovery Rate (FDR) ≥ 0.05. **e**, Correlation of the RNA expression and promoter accessibility in post-confinement cells. **f**, Genome browser tracks of ATAC-seq signal at representative DEGs across different confinement conditions. Gene locus annotated with UCSC browser style representation.

Comparison of differentially expressed genes revealed minimal overlap among conditions, indicating that the majority of transcriptional changes are specific to each confinement level (**Fig. 5b, Extended Data Fig. 7b-c**). This confinement-specific signature was further supported by expression patterns of shared DEGs, which varied between conditions as shown in the heatmap (**Fig. 5c**).

To gain a broad view of the biological processes associated with confinement, we performed gene set enrichment analysis (GSEA) using the Gene Ontology: Biological Process (GO:BP) database, which offers more granular and functionally resolved annotations and is therefore particularly useful for early-stage profiling of largely uncharacterized phenomena such as confinement-induced state changes. GSEA revealed that migration through low confinement was primarily associated with biological processes related to cytoskeletal remodeling and cell-ECM interactions, while migration through high confinement was associated with a broader signature of cell stress and chromatin regulation (**Fig. 5d**). Notably, biological processes enriched specifically under high confinement included wound healing, cell spreading, platelet aggregation, homotypic cell-cell adhesion, and cellular response to unfolded proteins, highlighting both a mechanically-induced stress response and morphological remodeling. In contrast, both wide control channels and low confinement conditions were associated with increased biosynthetic and translational activity. Interestingly, migration through low confinement also showed specific enrichment in regulation of protein localization to Cajal bodies, which may provide a mechanistic basis for our previous observation of phase-separating condensate formation under similar specific conditions^53^.

To further dissect the regulatory mechanisms underlying these transcriptomic changes, we performed ATAC-seq on matched conditions. We compared fold changes in chromatin accessibility at promoter regions with the corresponding expression fold changes from RNA-seq for all genes with significant expression differences (adjusted p value ≤ 0.05), which revealed a global reduction in promoter accessibility in high confinement channels, while low confinement did not cause a major change in accessibility (**Fig. 5e**). Consistent with confinement-induced changes in chromatin organization, high confinement also significantly reduced the coefficient of variation of nuclear DAPI intensity, indicating a more homogeneous spatial distribution of nuclear chromatin (**Extended Fig. 7f-i**). Genome browser views showed examples of confinement-induced decreases in accessibility at select loci (**Fig. 5f**). Representative genes are involved in stress adaptation and key regulatory pathways in cancer progression. For example, the PDCD6 gene codes for a calcium-binding protein that has been implicated in apoptosis^54,55^, the FAM20c gene produces a key protein kinase that promotes the invasion and metastasis of multiple cancers^56^, and the GNG2 gene acts as a tumor suppressor in breast cancer that also inhibits brain metastases^57,58^. Interestingly, the wide control channels also exhibited a global reduction in chromatin accessibility, further highlighting the distinct adaptations that occur in unconfined 3D spaces compared to standard 2D substrates. One possible explanation is that despite the absence of strong physical confinement, cells in wide control channels are experiencing a 3D microenvironment in which they interact with the channel walls and adopt morphologies that are distinct from those observed on flat 2D substrates. (**Fig. 3a)**. Previous work has shown that negatively curved adhesive environments can alter cell morphology, cytoskeletal organization, and adhesion distribution, suggesting that curvature-associated adhesion remodeling may contribute to the chromatin accessibility changes observed in wide channels^59^.

### Confinement induces cancer-associated and stress-responsive transcriptional programs

Cancer cell migration has been shown to correlate with metastatic potential and thus is an important factor in clinical prognosis^18,60^. Our microchannel platform allows us to specifically isolate and examine cells that have successfully traversed extreme confinement, making it possible to identify molecular differences associated with migration through increasingly confining physical environments. We first characterized these responses in HT1080 fibrosarcoma cells before extending the analysis to MDA-MB-231 breast cancer cells to determine which transcriptional responses were conserved across distinct cancer backgrounds.

Starting with high confinement-induced DEGs (high vs 2D), we compared their expression levels directly to the low confinement condition using significance thresholds alone. From this pool, we identified four groups of genes based on their confinement-specific transcriptional regulation modalities (**Fig. 6a-e**). Genes in Group 1 respond specifically to high confinement (i.e. are unchanged across 2D and low confinement) (**Fig. 6b**), while those in Group 2 exhibit a gradual, confinement dose-dependent pattern that changes significantly from 2D to low confinement to high confinement (**Fig. 6c**). Genes in Group 3 respond to confinement in general, with both low and high confinement eliciting a differential response but not compared to each other (**Fig. 6d**). Finally, Group 4 genes (**Fig. 6e**) show a differential response at high confinement, but low confinement does not elicit a significant response compared to either high confinement or 2D, suggesting more complex confinement-dependent regulation that could not be readily assigned to the other response patterns.

**Fig. 6.**
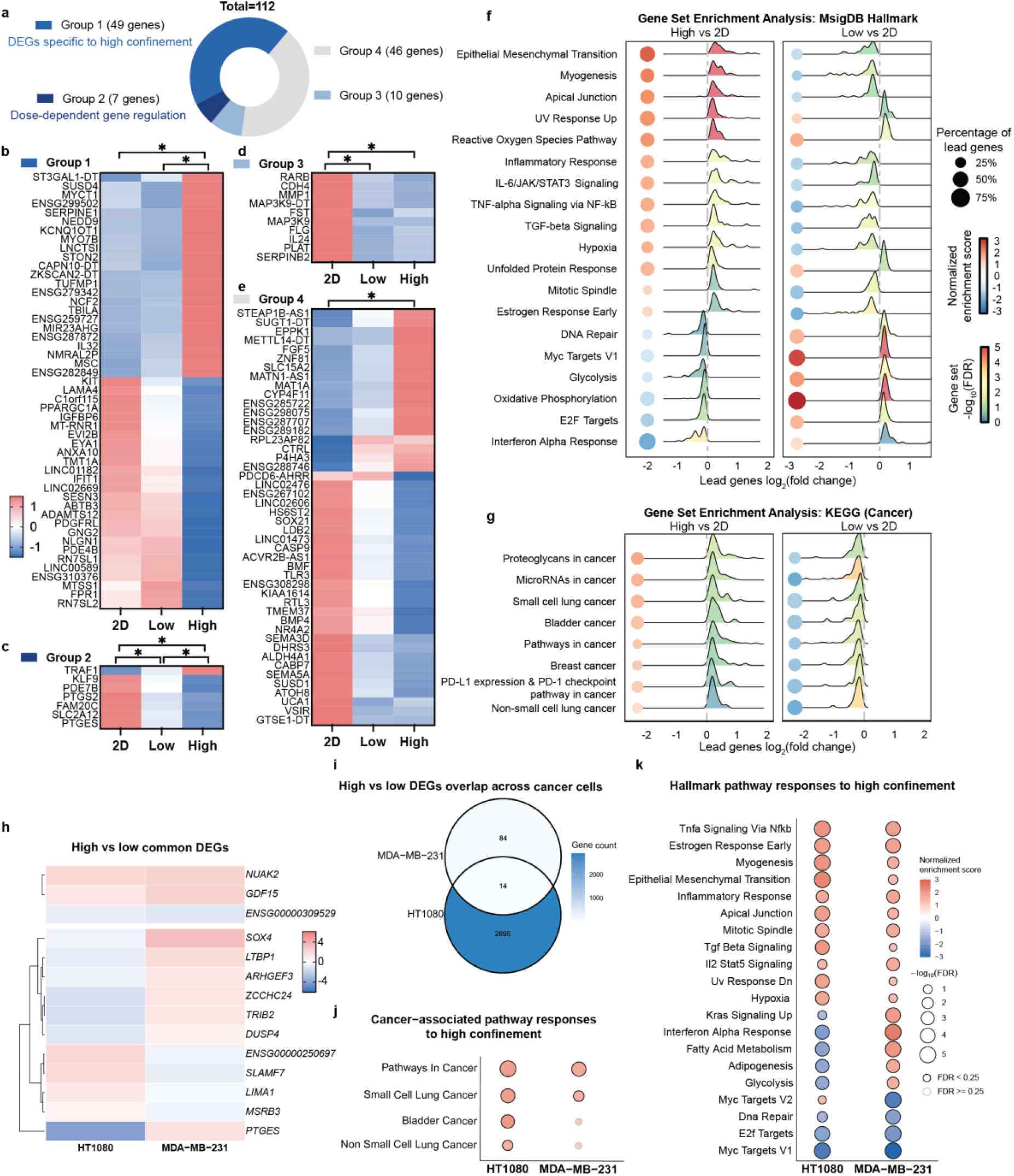
| Confinement-dependent transcriptomic reprogramming in cancer cells. **a**-**e**, DEGs from high confinement vs 2D comparison in HT1080 cell were classified into four groups based on expression trends across 2D, low-confinement, and high-confinement conditions. Heatmaps show z-scored gene expression for Group 1 (**b**), Group 2 (**c**), Group 3 (**d**), and Group 4 (**e**). * indicates adjusted p value ≤ 0.05 from gene expression differential analysis between each pair of conditions; ns, not significant. **f**-**g**, GSEA analysis of MSigDB hallmark pathways and KEGG cancer-related pathways. Enriched pathways are shown for high confinement vs 2D and low confinement vs 2D. **h**, Heatmap showing z-scored expression of common DEGs between HT1080 and MDA-MB-231 cells in the high vs low confinement comparison. **i**, Venn diagram of DEGs between HT1080 and MDA-MB-231 cells in the high vs low confinement comparison. DEGs were defined by adjusted p value < 0.05 and baseMean ≥ 10, irrespective of the direction of expression change. **j**-**k**, GSEA analysis of KEGG cancer-associated pathways (**j**) and MSigDB Hallmark pathways (**k**) in HT1080 and MDA-MB-231 cells under high vs low confinement. Dot color indicates NES, dot size indicates −log10(FDR), and outlines indicate FDR < 0.25 or FDR *≥ 0.25*.

Among the high confinement-responsive genes in HT1080 cells, we identified several cancer-associated genes that were strongly induced following migration through the 30 µm² channels. These included NEDD9, a focal adhesion scaffolding protein associated with enhanced migration and epithelial-to-mesenchymal transition^61^, SERPINE1, a regulator linked to invasive and stress-adapted cancer phenotypes^62–64^, CCN2, which is associated with extracellular matrix remodeling^65–67^, and CD274, which encodes the immune checkpoint protein PD-L1^68,69^. Protein expression of these targets was subsequently examined by immunofluorescence. PD-L1 showed the clearest agreement with the RNA-seq data, with increased protein abundance under high confinement while remaining lower or largely unchanged under the other channel conditions (**Extended Data Fig. 8a**). SERPINE1 showed a similar confinement-dependent pattern and was significantly higher under high confinement than in the wider channels, although the increase relative to 2D was not significant (**Extended Data Fig. 8b**). NEDD9 protein abundance was also higher under high confinement than in the wider channel conditions, although the increase relative to 2D observed at the transcript level was not reproduced (**Extended Data Fig. 8c**). In contrast, CCN2 protein abundance increased across all channel conditions and did not recapitulate the confinement-specific transcriptional response (**Extended Data Fig. 8d**). These results show that several high-confinement-responsive transcriptional changes are reflected at the protein level, while also highlighting gene-specific differences between transcript and protein abundance. Integrated transcription factor inference further identified increased activity of SMAD2/3- and SNAI1-associated regulatory networks under high confinement. Consistent with this response, SERPINE1, a downstream target of these pathways, was among the most strongly upregulated genes in the dataset (**Extended Data Fig. 8e-f**).

We then compared transcriptomes from high confinement and 2D controls, as well as from low confinement and 2D controls, focusing on pathways that showed opposite enrichment patterns or were selectively enriched in high confinement (**Fig. 6f**). Notably, migration through highly confining microchannels led to upregulation of hallmark pathways linked to the epithelial-to-mesenchymal transition, reactive oxygen species pathways, inflammatory response, and TNFα/NF-κB signaling, all of which are commonly associated with aggressive and stress-adapted cancer phenotypes. Many of these pathways were unchanged or suppressed under low confinement, further underscoring the distinct state induced as a consequence of migration through extreme confinement. Examining cancer-associated KEGG pathways, GSEA analysis revealed significant enrichment of pathways implicated in tumor progression, including proteoglycans in cancer, microRNAs in cancer, as well as certain specific types of cancers such as small cell lung cancer, exclusively in high-confinement conditions (**Fig. 6g**). These results indicate that severe confinement induces a distinct cancer-associated transcriptional state in HT1080 cells.

To determine whether these responses extended beyond HT1080 cells, we performed RNA-seq on MDA-MB-231 cells following migration through the same confinement conditions. MDA-MB-231 cells exhibited a largely distinct transcriptional and pathway response, although some responses were shared with HT1080 cells, including reactive oxygen species signaling (**Extended Data Fig. 9**). We next directly compared high and low confinement across both cancer cell lines to identify a shared transcriptional signature of severe confinement. This analysis identified 14 genes that were significantly altered in both HT1080 and MDA-MB-231 cells (**Fig. 6h and i**). Among these shared genes, NUAK2 and GDF15 were consistently upregulated under high confinement in both cell lines, although other genes showed cell-type-specific differences in direction or magnitude (**Fig. 6h**). At the pathway level, the two cancer models showed further convergence in cancer- and stress-associated programs, including epithelial-to-mesenchymal transition, inflammatory response, TGF-β signaling, and TNFα/NF-κB signaling (**Fig. 6j-k**). We further identified six genes whose expression progressively scaled with increasing confinement across both cancer cell lines (**Extended Data Fig. 8g**). Four showed the same progressive downregulation in both cell types, while two showed opposite trends between the two cell types (**Extended Data Fig. 8h**). These responses reveal a confinement-responsive program of cancer cell adaptation that is conserved across distinct cancer backgrounds, one that is generally encoded transcriptionally as a graded cellular response to increasing levels of confinement.

## Discussion

Nearly every cell in the body experiences confinement; thus, building tools that mimic this confinement is of great importance for understanding, mitigating, and leveraging confinement-sensitive cell behavior. While great strides have been made in our understanding of mechanosensitive cellular responses to confinement^44,70–72^ and even led to the development of accurate diagnostic approaches^73^, a critical gap in the confinement toolbox is a method that allows for confined migration with efficient recovery of post-migratory cells. Excluding tools that use non-physiological flow to push non-adherent cells through constrictions^17,74^, the current throughput for post-confined migration cells is roughly 150 cells per well in a six well plate^33^. While a very recent approach has utilized microfluidics to yield approximately 200 cells per device, this system requires the use of potentially confounding non-physiological flow^32^. Here, we have gone orders of magnitude past these benchmarks, obtaining 100,000 cells per square centimeter membrane, with a theoretical maximum of 1,200,000 cells per standard six well plate. This system is design-flexible, allowing for a wide range of confining cross-sectional areas, lengths, geometries, and interconnectivity.

Importantly, and underscored by our observation here of confined migration by multiple diverse cell types (**Figure 4)**, this method is easily adaptable by a wide variety of bioengineering and cell biology labs. Any lab with expertise in microfabrication will be able to replicate this method and build their own high-content microchannel membrane system for their own unique biological focus. This is especially important given the well-known challenges inherent to producing high-aspect ratio structures in the Z-axis. With the inclusion of 3D printed resin inserts, labs equipped for standard cell biology will be able to utilize this system the same way they currently utilize standard commercial transwell assays. To facilitate this, we have assembled inserts allowing for culture in specific formats depending on the usage case: 24 well plates for image-based assays that prioritize efficiency and decreased reagent volume, and 12 well plates for lysis-based assays that prioritize lysate production.

Despite these advantages, the current fabrication workflow remains more time-intensive than the use of commercially available confinement platforms. Device production requires multiple rounds of PDMS casting, plasma bonding, stacking, and vibratome sectioning over the course of several days. In particular, the vibratome slicing step requires careful optimization of cutting parameters, as the elastic nature of PDMS can result in incomplete sectioning or membrane deformation at higher blade feed rates. Furthermore, while the membrane format is well suited for endpoint imaging and visualization of subcellular features, the increased working distance required to image through the device limits compatibility with high numerical aperture objectives and super-resolution imaging modalities. Live-cell migration can also be visualized and migration speed quantified, although the vertical channel orientation requires repeated z-stack acquisition, making conventional horizontal PDMS-on-glass microchannels better suited for detailed or high-throughput analysis of single-cell migration dynamics.

Importantly, these considerations do not diminish the primary advantage of the platform, namely its compatibility with large-scale downstream analyses. This is evident in our results, where we obtained transcriptional and epigenetic signatures of confined migration via RNAseq and ATACseq. Despite the rising accessibility of advanced omics techniques, no work has directly examined how self-directed cell migration through defined levels of physiological confinement influences gene expression and chromatin accessibility, largely because conventional confinement platforms have not enabled sufficient recovery of post-confined cells for robust sequencing-based analysis. While single-cell omics approaches are increasingly powerful for resolving cellular heterogeneity, bulk RNAseq and ATACseq remain valuable for defining robust population-level signatures and identifying molecular responses to confinement. These sequencing approaches revealed some expected outputs, including upregulation of genes involved in the epithelial-to-mesenchymal transition and a wide variety of cancer-associated pathways, as well as several unexpected and exciting outputs, including a strong transcriptional and epigenetic distinction between extremely wide control channels and intermediate channels that allow the cell to engage in three dimensions but do not cause nuclear deformation. This particular finding suggests that on 2D surfaces, where traction force buildup resulting from strong apicobasal polarity can cause significant nuclear deformation, unconfined cells are perhaps more similar to cells under extreme confinement than was previously thought. Thus, the transcriptional and epigenetic differences between low confinement and high confinement may be much more informative for understanding physiological responses to confinement than on 2D control surfaces. Moreover, the ability to collect large numbers of post-confined cells may also enable future single-cell omics approaches, where sufficient starting material remains important for sample recovery, library preparation, and quality control.

Beyond the compatibility with lysis-based assays, which we have demonstrated here, this system is also compatible with the collection of intact post-confinement cells. After allowing cells to migrate to the exit surface of the membrane, it is straightforward to harvest cells either by enzymatic or mechanical detachment. Again, despite the biological importance of confinement in regulating cell behavior, there are no current methods that allow for the bulk collection of tens of thousands of cells after exposure to specific levels of confinement. This approach has a number of important applications in the future, including the isolation of interesting and/or clinically relevant subpopulations of cells, e.g. the collection of the most invasive cancer cells within a heterogeneous patient sample. The inverse is also true; as mechanical adaptation of cells can lead to persistent changes in their cell state^4,16,75^ or behavior, utilizing a bulk confinement system like this will allow researchers to collect populations of mechanically-pretreated cells for downstream clinical use^74^ with exciting possibilities like stem cells for regenerative medicine or immune cells for cancer immunotherapy. Beyond this, as serial passaging of existing cells has long been used to produce new cell lines with specific functions^76,77^, this method can also be quickly adapted to yield new mechanically-adapted cell lines for further fundamental research on the molecular underpinnings of confined migration. Finally, similar to transwell assays, if a cell emerges from confinement and does not remain attached to the exit surface of the membrane, it will fall to the base of the well. This allows for straightforward separation of subpopulations of cancer cells that A) do not migrate, B) migrate but remain adherent, and C) migrate and undergo an adhesion-to-suspension transition (AST). This AST phenomenon has only recently been focused on as a key driver of metastasis^78^, meaning that the method presented here would be highly useful for further fundamental research on this cell behavior, as well as any other modality in which cells lose adhesion to the substrate.

Overall, as the field of confinement mechanobiology continues to explore how physical forces shape cellular identity and function, our system provides a versatile and scalable framework to translate these insights into experimentally tractable models, with the potential to reveal new targets and mechanisms relevant to both health and disease.

## Methods

### Photolithography

Rectangular microchannels with heights of 5 µm (high confinement, ‘high’), 10 µm (low confinement, ‘low’), and 20 µm (no confinement, ‘wide’) were fabricated using silicon wafer master molds. For periodic confinement channels, microchannels with 10 µm x 10 µm (H x W) cross-sections were fabricated with periodic 15 µm long constrictions (10 µm x 3 µm cross-sections) spaced at 50 µm intervals along the channel length (**Extended Data Fig. 4c-e**).

Prior to photoresist application, silicon wafers were dehydrated by baking at 180-200 °C for fifteen minutes. Photoresists (SU-8 3005 for 5 µm thickness, SU-8 3010 for 10 µm thickness, and SU-8 3025 for 20 µm thickness; Kayaku Advanced Materials, MA, USA) were spin-coated at 4200 rpm (5 µm), 2600 rpm (10 µm), and 4000 rpm (20 µm) for 30 seconds. A two-step soft bake process was then applied to each wafer: one minute at room temperature followed by four minutes (5 µm), eight minutes (10 µm), or ten minutes (20 µm) at 95 °C. The 5 µm and 10 µm photoresist layers were exposed to 130 mJ/cm² ultraviolet light (365 nm wavelength, SUSS Microtec MJB4 mask aligner) through a custom soda-lime optical mask. The 20 µm photoresist layer was exposed to 700 mJ/cm² ultraviolet light (375 nm wavelength, MLA 150 Maskless Aligner; Heidelberg Instruments, Germany) using the same mask type. All samples underwent a post-exposure bake: one minute at 65 °C followed by two minutes at 95 °C (5 µm and 10 µm layers) or four minutes at 95 °C (20 µm layer). The wafers were developed in SU-8 developer (Kayaku Advanced Materials, MA, USA) at room temperature, rinsed with isopropyl alcohol, and dried with nitrogen gas. A final hard bake was performed at 130 °C for five minutes, followed by sequential cooling steps at 95 °C and 65 °C (one minute each).

### Membrane fabrication

PDMS was mixed with a 1:10 curing agent to base ratio (SYLGARD 184 Silicone Elastomer) and cast over the silicon master mold. For handling stability, the first layer was approximately 500 µm thick, produced by spin-coating eight grams of uncured PDMS mixture onto the wafer at 300 RPM for thirty seconds. To create the subsequent thin 100 µm PDMS layers, five grams of uncured PDMS was spin-coated onto the wafers at 1000 RPM for thirty seconds. All samples then underwent vacuum degassing at 10 mbar for ten minutes to remove bubbles, followed by curing at 80 °C for one hour. As thin PDMS films are prone to tearing, they were gently peeled from the wafer using gloved fingers rather than forceps. For trimming, thin PDMS films were first placed onto a thicker PDMS support block before the square patterned regions were cut to size using a razor blade. The patterned surface of each PDMS film was tracked throughout assembly to ensure that the microchannel-containing side was correctly oriented for attachment to the next PDMS layer.

PDMS film attachment between layers was achieved via three-minute oxygen plasma activation of both surfaces (Harrick Plasma Cleaner). Activated PDMS films were slowly attached with gentle pressure to prevent the formation of bubbles, while taking care to align the patterned regions by eye. The assembled layers were then baked at 80 °C for twenty minutes to strengthen covalent bonding between PDMS films.

After eight to nine thin PDMS films had been attached to form a multilayer block, multiple blocks were joined to increase the total number of channel layers. A thin layer of uncured PDMS was applied only to the interface between multilayer blocks to strengthen inter-block bonding. As uncured PDMS applied between multilayer blocks fills the channel layer directly adjacent to the bonding interface, this interfacial layer was treated as sacrificial, and care was taken to minimize spreading into neighboring channel layers. The combined PDMS stack underwent vacuum degassing at 10 mbar for ten minutes to remove bubbles, followed by curing at 80 °C for one hour. Stacked PDMS blocks were pre-cut into approximately 1 cm x 0.6 cm x 0.6 cm cubes for optimal vibratome slicing performance. During this step, the outer edges of the stacked blocks were removed to exclude regions where excess uncured PDMS may have spilled over during block assembly.

A Leica VT1200 S automated vibrating blade microtome was used to slice the PDMS blocks. The vibratome blade feed rate was set at 0.02-0.03 mm/s, with vibration amplitude at 1 mm and a slicing thickness of 150 µm. Slower blade feed rates were required to fully section the elastic PDMS block and minimize membrane deformation during cutting. Sliced PDMS membranes were stored in PBS solution at 4 °C for future use. Membranes were inspected after slicing, and samples with visible tearing, incomplete sectioning, severe deformation, poor layer bonding, or obvious channel blockage were excluded from downstream experiments.

### Cell culture insert fabrication

Cell culture inserts were fabricated using a stereolithography (SLA) 3D printer (Original Prusa SL1S SPEED) with a biocompatible photopolymer resin (Prusament Resin Model Alabaster White 1kg). Insert designs were created in Onshape and exported as standard tessellation language (STL) files. Printing parameters included a layer thickness of 0.025 mm, an exposure time of two seconds/layer, and an initial exposure time of 20 seconds, optimized for structural fidelity. Post-processing involved rinsing in 100% Isopropyl Alcohol to remove any residual resin, followed by curing at 405 nm for seven minutes to ensure complete polymerization, all of which was done in an Original Prusa Curing and Washing Machine (CW1S).

Printed resin inserts were sonicated in DI water for 30 minutes at 45 °C and washed in 100% ethanol to remove residual uncured resin. For immunofluorescence samples, PDMS membranes were attached to 24 well plate resin inserts by low viscosity silicone (Wirosil Dublier-Silikon) to enable straightforward detachment post-staining. For sequencing samples, PDMS membranes were attached to 12 well plate resin inserts by uncured PDMS (SYLGARD Silicone Elastomer) and then baked at 80 °C for twenty minutes.

Membrane-attached inserts were exposed to oxygen plasma (Harrick Plasma Cleaner) for thirty seconds and then coated with rat tail collagen I (GIBCO) dissolved in 0.1% acetic acid solution at a concentration of 15 µg/mL. Coated membrane-attached inserts were UV sterilized before cell culture experiments. For comparisons with microchannel systems, Corning® Transwell® 24 well plates (5.0 μm pore size, polycarbonate membrane) were purchased.

### Cell culture

The HT1080 cell line was a kind gift from Prof. Alexander Bershadsky’s lab. HT1080, MDA-MB-231, and HFF-1 cells were cultured at 37 °C with 5% CO2 and passaged when cell confluency reached 80% with Trypsin-EDTA (0.05%, no phenol red, GIBCO). Complete culture media consisted of 10% Fetal Bovine Serum (GIBCO), 1% Penicillin-Streptomycin (GIBCO), and 89% Dulbecco’s Modified Eagle Medium (DMEM) (GIBCO). Cell cultures were tested for Mycoplasma contamination routinely with the Mycolor One-Step Mycoplasma Detector (Vazyme, D201).

Human bone marrow-derived mesenchymal stem cells (hBM-MSC) were purchased from ATCC. Cells were maintained in low-glucose DMEM media (GIBCO) with 10% Fetal Bovine Serum (GIBCO) and 1% Penicillin-Streptomycin (GIBCO). hBM-MSCs were cultured at 37 °C with 5% CO2 and maintained under 70% confluency up until passage 6.

To obtain primary bone marrow-derived macrophages, bone marrow was retrieved from the femurs and tibias of 4-week-old female C57BL/6 mice under the Tissue Sharing Program of the National University of Singapore. Euthanasia was performed via CO2 asphyxiation by Institutional Animal Care and Use Committees (IACUC)-approved personnel, and all animal procedures were performed in accordance with IACUC protocols. Femur and tibia exposure was done through the dissection of mouse lower limbs. Briefly, cuts were made at the knee, ankle, and hip joints to obtain the necessary bone pieces. Connective tissue was manually removed, and the bones were transferred to 0.6 mL Eppendorf tubes. Centrifugation at 16,000 RPM for ten seconds was performed to isolate the bone marrow, followed by red blood cell (RBC) lysis using 1X RBC Lysis Buffer (eBioscience™ Catalog No. 00-4333-57) for 10 minutes at room temperature. A second round of centrifugation at 1,000 RPM for 3 minutes was then performed, with the cell pellet subsequently transferred to cell culture flasks for culture and expansion. Harvested cells were maintained in 10% Fetal Bovine Serum, 1% Penicillin-Streptomycin, and RPMI 1640 media (Thermofisher), supplemented with 20 ng/mL of macrophage colony stimulating factor (MCSF, Sigma Aldrich M6518-10UG) for at least seven days before usage to ensure complete differentiation into macrophages.

### Fluorescent staining

Transwell pore and microchannel integrity post-slicing were visualized with a Dextran-Alexa Fluor™ 594 (125 µg/ml, 10000 MW, Anionic, Fixable, ThermoFisher) solution in 0.1% acetic acid. Collagen coating on PDMS was assessed via labelling with CNA35-eGFP (a kind gift from Prof. Jennifer Young).

Cells were fixed with 4% paraformaldehyde (PFA) in PBS (Santa Cruz, CAS 30525-89-4) at room temperature, then permeabilized and blocked with 0.1% Triton X-100 and 2.5% bovine serum albumin (BSA) in PBS at room temperature for 30 minutes. For cellular and nuclear morphology characterization in fixed cells, primary and secondary antibodies were dissolved in 1% BSA solution in 0.1% Tween-20. Cells were incubated with a primary antibody for Paxillin (1/500, ab32084), Phospho-Myosin Light Chain 2 (1/400, mAb #3675), SERPINE1 (0.1µg/mL, ab222754), NEDD9 (1:200, ab18056), CCN2 (1:100, ab6992) and CD274 (2µg/mL, ab205921) overnight at 4 °C, followed by three washes with 0.1% PBST. Secondary antibodies (Donkey anti-Mouse IgG (H+L) Highly Cross-Adsorbed Secondary Antibody, Alexa Fluor™ Plus 488, 1/2,000; Donkey anti-Rabbit IgG (H+L) Highly Cross-Adsorbed Secondary Antibody, Alexa Fluor™ 647, 1/2,000), Alexa Fluor 568 Phalloidin (1/400, A12380) and Hoechst 33342 (1/1,000, H1399) were incubated for four hours at room temperature. Immunofluorescence validation was performed using conventional PDMS-on-glass microchannel devices following fabrication and immunofluorescence protocols previously established in our laboratory^10,16,50^.

### Live migration tracking

Two adhesive, 250 µm thick spacers (iSpacer) were affixed to a 35 mm glass bottom dish (Cellvis) 3 mm apart. PDMS membranes were dried and attached over the spacers, ensuring that membranes were taut over the gap between spacers. The assembled membranes were exposed to oxygen plasma and coated with collagen I as described previously. The assembly was then washed with PBS and sterilized using UV exposure. Prior to cell seeding, PBS in the assembly was replaced with cell culture medium and equilibrated in the incubator. Medium was aspirated from the dish as well as from the top of the membrane but maintained within the gap between spacers. HT1080 NLS-GFP LifeAct-tdTomato cells were seeded on the top of the membrane at a density of 2 million cells/ml to fully cover the membrane and allowed to attach briefly before the assembly was fully immersed in cell culture medium. Live imaging was performed on a Yokogawa W1 spinning disk confocal microscope using a 40X LWD objective with 1 µm step size. Z-stacks were acquired every 10 minutes, and timelapse data tracking the nucleus of the cells was analyzed using Fiji.

### RNA and ATAC sequencing

Bulk RNA-seq samples were extracted with a Monarch® Total RNA Miniprep Kit (#T2010S) following the product protocol for cultured mammalian cells. Cell lysates from each biological replicate were collected and frozen at -80 °C before RNA extraction. On-column DNase I treatment was performed for enzymatic removal of residual gDNA. RNA concentration was quantified using a NanoDrop 8000 Spectrophotometer (Thermo Fisher Scientific). ATAC sequencing samples were collected and extracted with a Hyperactive ATAC-Seq Library Prep Kit for Illumina (Vazyme, #TD711) and index sequences were added to the sample with the TruePrep Index Kit V2 for Illumina (Vazyme, #TD202).

Standard bulk sequencing was performed on the NovaSeq X Plus sequencing platforms (NovogeneAIT Genomics Singapore Pte Ltd). RNA sequencing data was processed using the nf-core/differentialabundance pipeline, which includes quality control, annotations, and differential expression analysis using DESeq2. Gene set enrichment analysis (GSEA, https://www.gsea-msigdb.org/gsea/index.jsp) was performed in Python using the gene sets GO_Biological_Process_2025, MSigDB_Hallmark_2020, and KEGG_2021_Human. Computation of z-score heatmaps was performed in Python using custom scripts. Differentially expressed genes were identified by (|log2(FoldChange)|≥ 1, adjusted p-values ≤ 0.05). Final data visualization was carried out using R (ggplot2) and GraphPad Prism. Transcription factor activity was inferred from RNA-seq data using a regulon-based approach, in which coordinated expression changes among curated downstream target genes were used to estimate the relative activity of each transcription factor across confinement conditions.

### Microscopy, image processing and quantification

Z-stack images were acquired with an Olympus FV3000 NIR Confocal Laser Scanning Microscope with a 30× objective (Olympus Plan Super Apochromat 30× Silicon Immersion Objective, NA 1.05, WD 0.8 mm) and a 1 µm z step.

Scanning electron microscopy was used to characterize the cross-sectional features of the microchannel membranes. Clean and dry PDMS samples were secured on a metallic mount using adhesive carbon tape and were uniformly coated with a 10 nm thick layer of platinum using a sputter coater (JFC-1600 Auto Fine Coater, JEOL). The prepared samples were then imaged using a scanning electron microscope (JSM-6010LV, JEOL) at an accelerating voltage of 20 kV.

Image processing was mainly performed using FIJI (ImageJ). This included image cropping, rotation, re-slicing, application of lookup tables, brightness/contrast adjustment, and maximum intensity projections for z-stack images. Imaris (Bitplane) was used for 3D rendering and visualization of microchannel structures. For analysis of fluorescence intensity through the microchannels, longitudinal intensity profiles were measured from fluorescent dextran-filled channels in Fiji. Channel positions were aligned to the 150 µm channel length, and fluorescence profiles from individual channels were normalized before averaging within each channel geometry. Cell and nuclear morphology measurements were conducted using CellProfiler, applying custom pipelines to segment fluorescently labeled nuclei and cytoplasm. For quantification of F-actin distribution, phalloidin fluorescence intensity was measured along the migration axis of individual cells. Cell length was normalized from the trailing edge (0%) to the leading edge (100%), and fluorescence intensity was normalized to the mean intensity of each cell. F-actin enrichment at the rear (5–20%) and front (80–95%) was calculated relative to the central cell body (35–65%). Chromatin organization was further assessed from DAPI-stained nuclei by calculating the coefficient of variation (CV) of intranuclear DAPI intensity, defined as the standard deviation divided by the mean intensity within each nucleus, following the protocol described by Martin et al. (2021)^79^.

### Data availability

Raw RNA-seq and ATAC-seq data have been deposited in the NCBI Sequence Read Archive under BioProject accession PRJNA1278927. Data will be made publicly available upon publication.

### Code availability

Custom analysis scripts and image-processing pipelines used in this study are available from the corresponding author upon reasonable request.

## Acknowledgments

All microscopy was carried out at the Microscopy Core Facility, MBI, National University of Singapore. We thank Dr. Gianluca Grenci, Mona Suryana, and Foo Sebastian Junwei of the Nano and Micro Fabrication Core Facility of the Mechanobiology Institute (MBI) for their assistance with the fabrication of silicon wafers. Dr. Paramasivam Kathirvel and Hemang Raj SINGH from the High-Throughput Molecular Genetics Core Facility provided great suggestions and assistance in executing the omics studies. We thank Dr. Arikta Biswas, Ng Boon Heng, and members of the Confinement Mechanobiology Lab for generously offering their valuable time and expertise. The authors acknowledge the support of the National Research Foundation of Singapore under an NRF Fellowship to AWH (NRFF13-2021-0114). This research is also supported by the National Research Foundation, Singapore, under its Mid-sized Grant (NRF-MSG-2023-0001).

---

**Extended Data Fig. 1.**
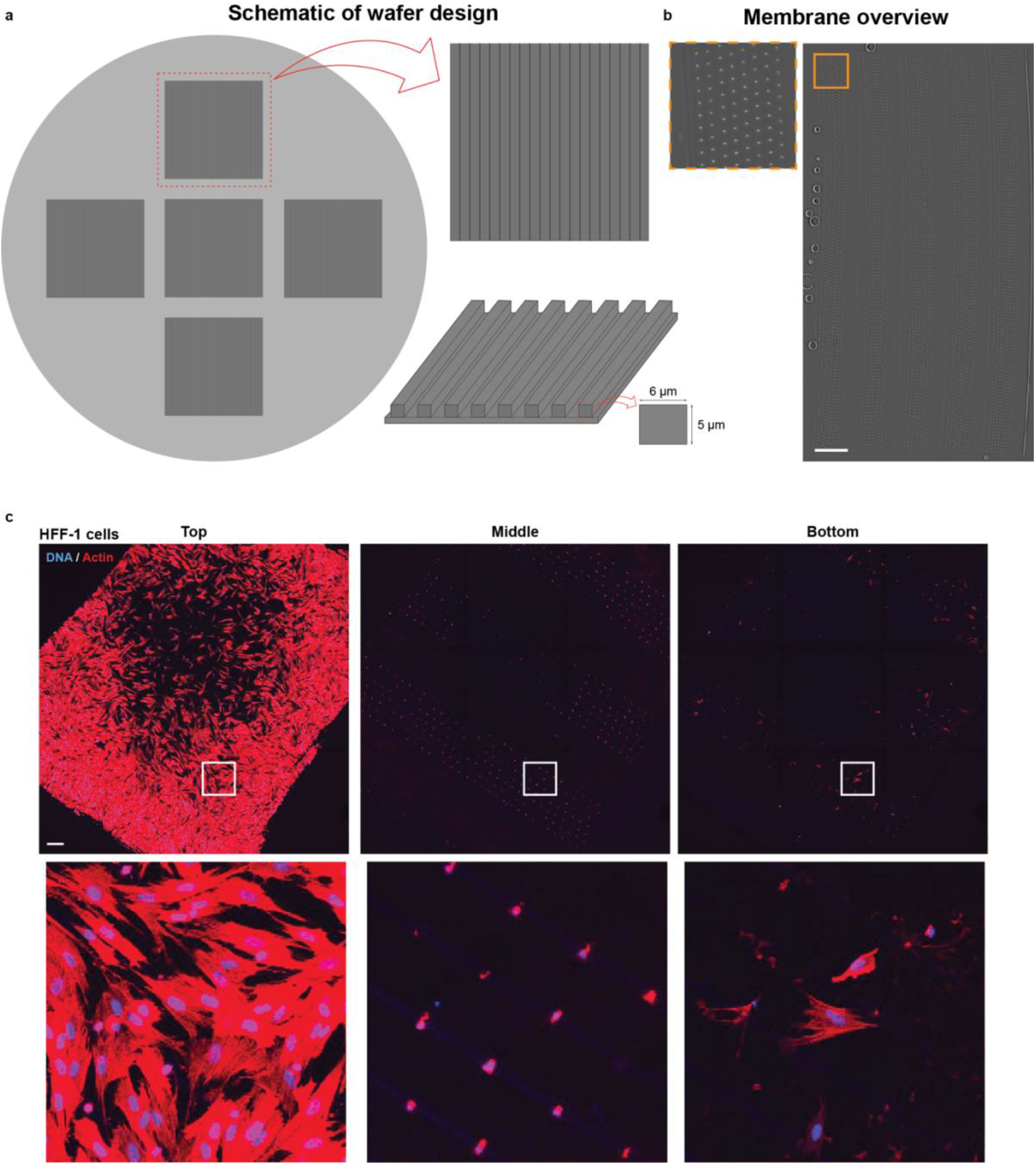
| Wafer design, channel density, and cellular coverage of the high-throughput microchannel membrane. **a**, Schematic of the photolithographic wafer layout used to fabricate the high-throughput microchannel membranes. Individual patterned regions contain arrays of long linear channel features with channel aspect ratios of 5 x 6 µm (high), 10 x 10 µm (low), and 20 x 20 µm (wide). **b**, Tiled phase-contrast image of a single high-throughput microchannel membrane, with inset highlighting the high density and uniform distribution of microchannels across the membrane. **c**, Representative tiled immunofluorescence images of HFF-1 cells seeded onto the high-throughput microchannel membrane. The top, middle, and bottom images illustrate the high number of cells that can simultaneously interact with and migrate through the microchannel array.

**Extended Data Fig. 2.**
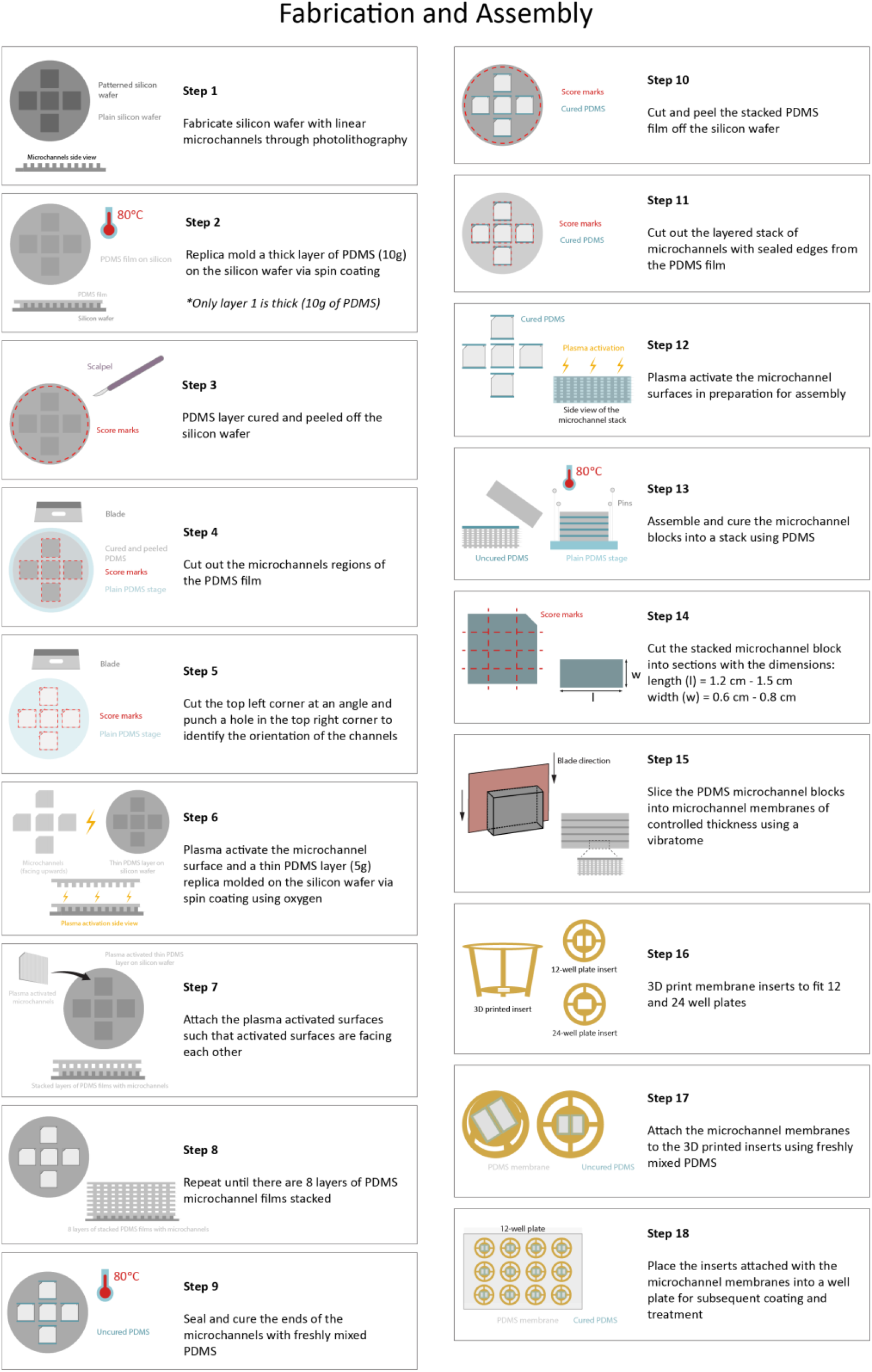
| Fabrication and assembly workflow for the high throughput microchannel membrane. Schematic illustrating the stepwise fabrication and assembly of the high throughput microchannel membrane. Linear microchannel features are first fabricated on a silicon wafer by photolithography and replicated in PDMS. Individual PDMS layers are cut, plasma activated, aligned, and sequentially bonded to form stacked microchannel blocks. The assembled blocks are then sectioned perpendicular to the channel axis using a vibratome to generate thin membranes containing arrays of microchannels. The resulting membranes are attached to custom 3D printed inserts using freshly mixed PDMS and placed into standard multiwell plates for subsequent coating and cell culture experiments.

**Extended Data Fig. 3.**
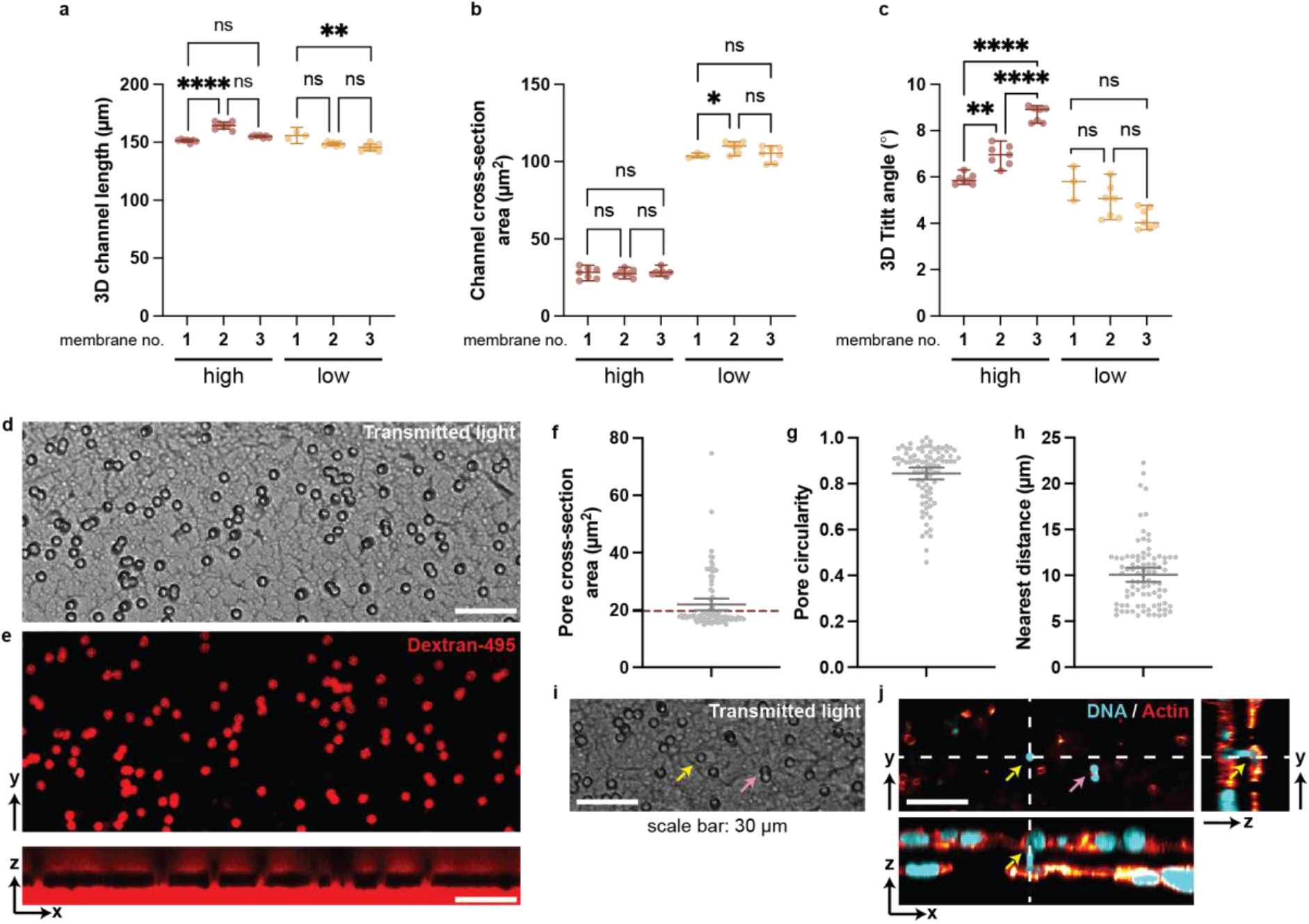
| Reproducibility of the high throughput microchannel membrane and limitations of commercial Transwell membranes for cellular confinement. a-c, Microchannel dimension measurements with three independent fabrication replicates presented separately: 3D microchannel length (a), channel cross-section area (b), and 3D tilt angle (c). Scatter plots: mean ± 95% C.I. d and e, Surface texture (d) and fluorescently labeled porous structure (e) of a conventional Transwell permeable membrane. f-h Variability in Transwell pore dimensions: cross-sectional area (f), circularity (g), and spatial distribution (h). Red dashed lines: expected pore size. Scatter plots: mean ± 95% C.I. i-j, Representative images of HT1080 cells penetrating Transwell pores. Due to limited membrane thickness, Transwell membranes fail to fully confine nuclei (yellow arrows), while irregular pore geometry deforms nuclei into corresponding shapes (pink arrows). Dashed lines: orthogonal planes. Scale bars: 30 µm.

**Extended Data Fig. 4.**
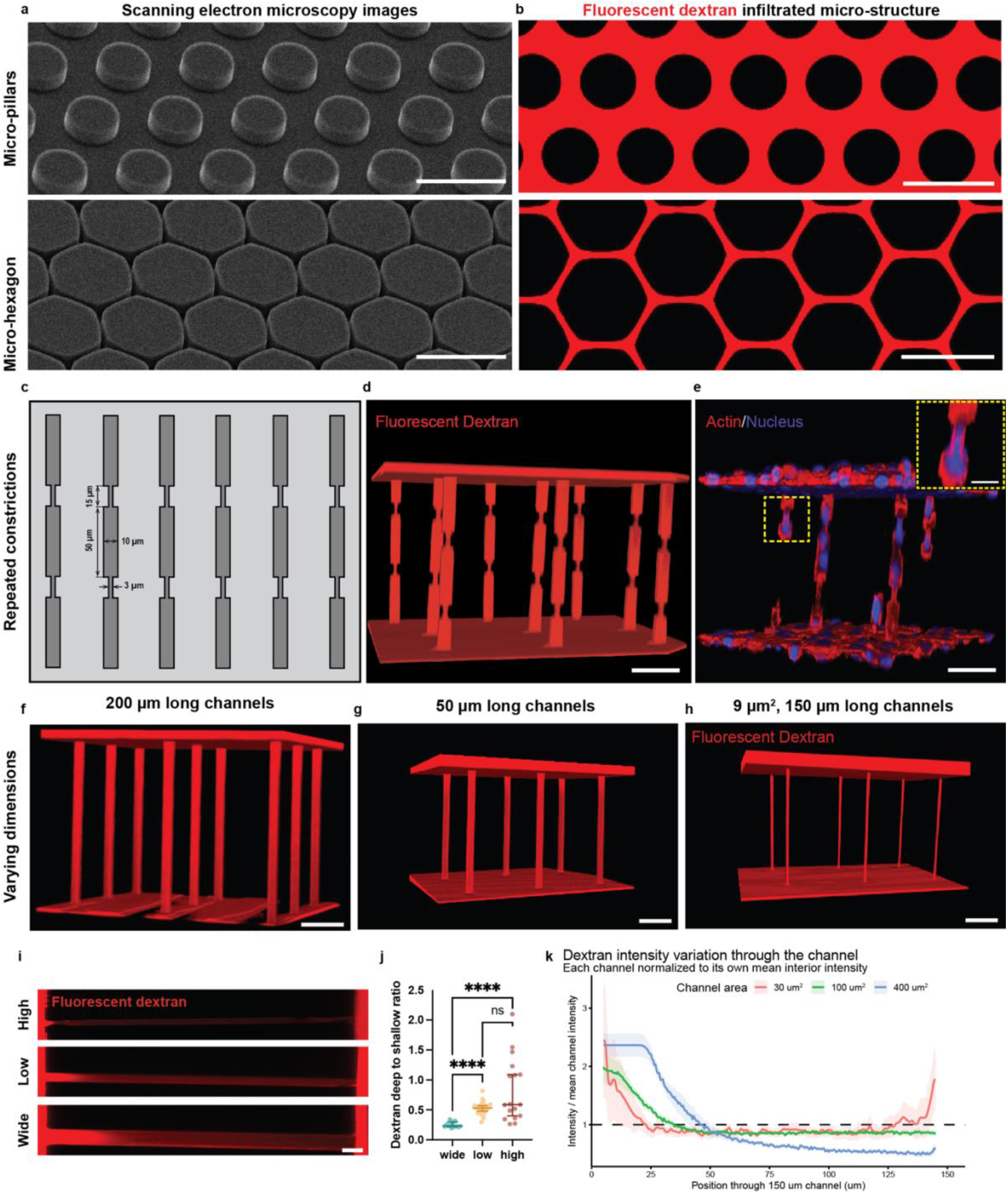
**| Versatile fabrication of confinement microstructures with diverse geometries. a-b**, Representative SEM images (left) and fluorescence images (right) of confining microstructures with inter-pillar (top) and hexagonal (bottom) paths. Scale bars: 100 μm. **c**, Schematic of repeated constriction designed with indicated dimensions. **d**, 3D reconstruction of fluorescent dextran-labeled repeated constriction within a membrane. Scale bar: 50 µm. **e**, 3D reconstruction of cytoskeletal and nuclear morphology of HT1080 cells migrating through repeated constrictions, with the boxed region magnified in the inset. **f-h**, 3D reconstructions of fluorescent dextran-labeled channels with varying dimensions: 200 µm long channels (**f**), 50 µm long channels (**g**), and 9 µm² cross-section and 150 µm long channels (**h**). **i**, Representative images of dextran-filled microchannels. Scale bars: 10 µm. **j-k**, Quantification of the ratio of fluorescent dextran intensity measured deep within the channel to that measured near the channel opening (**j**), and fluorescent dextran intensity profiles measured along 150 µm channels (**k**), showing a reduction of detected fluorescence within the channels, due to optical attenuation when imaging through the microchannel.

**Extended Data Fig. 5.**
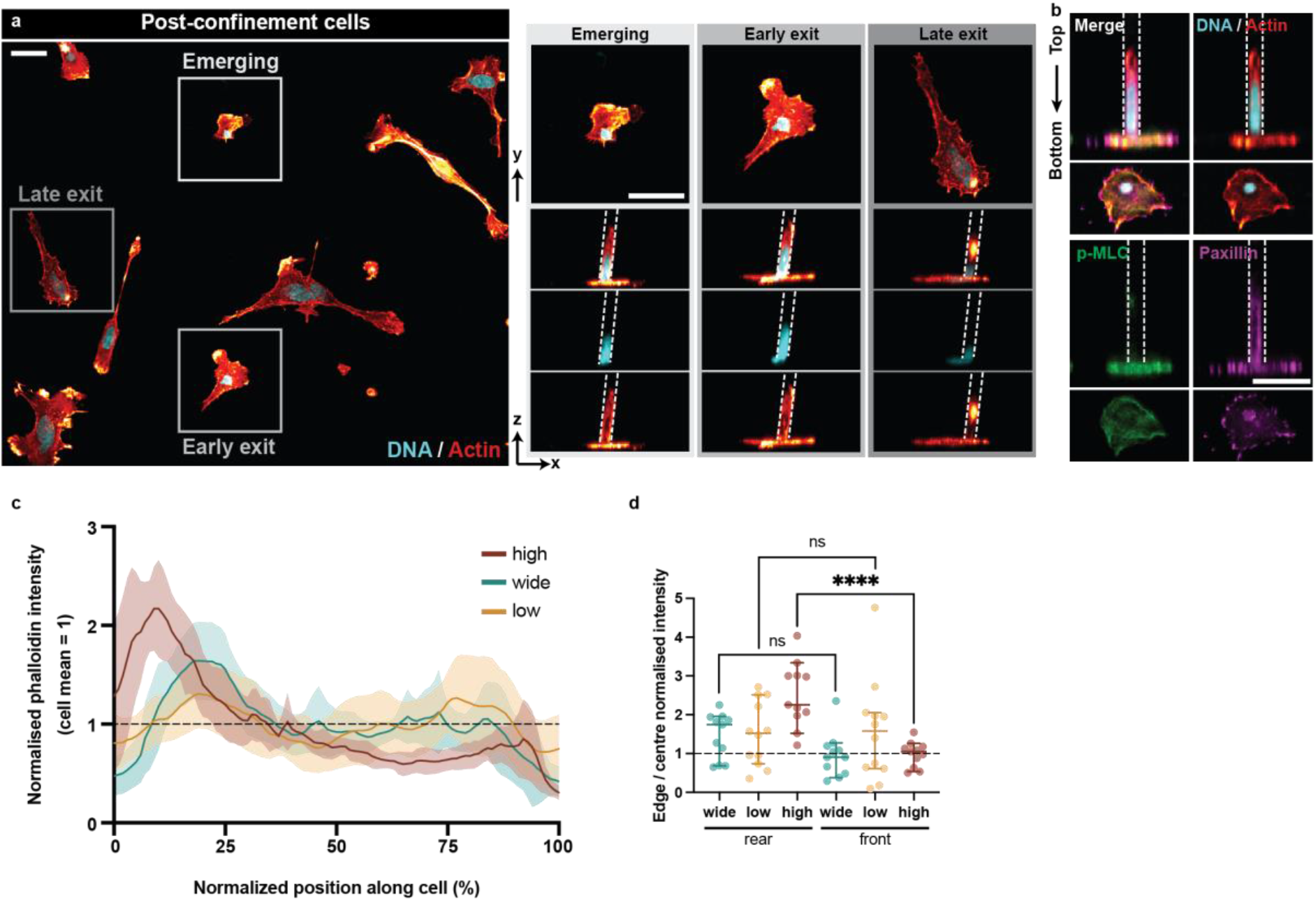
| Active cell emergence from confinement. **a**, Representative images of post-confinement cells at different stages of emergence, including cells just emerging, partially exited, and fully exited. **b**, Immunofluorescence staining (right) reveals enrichment of phosphorylated myosin light chain (p-MLC) and paxillin at the leading edge during confinement exit. Scale bars: 30 μm. Dashed lines: channel boundaries. **c**, Normalized actin intensity profiles along the cell axis of cells migrating through wide, low, and high confinement channels. **d**, Quantification of actin enrichment at the rear and front of cells under each confinement condition, showing increased rear cortical actin enrichment under high confinement.

**Extended Data Fig. 6.**
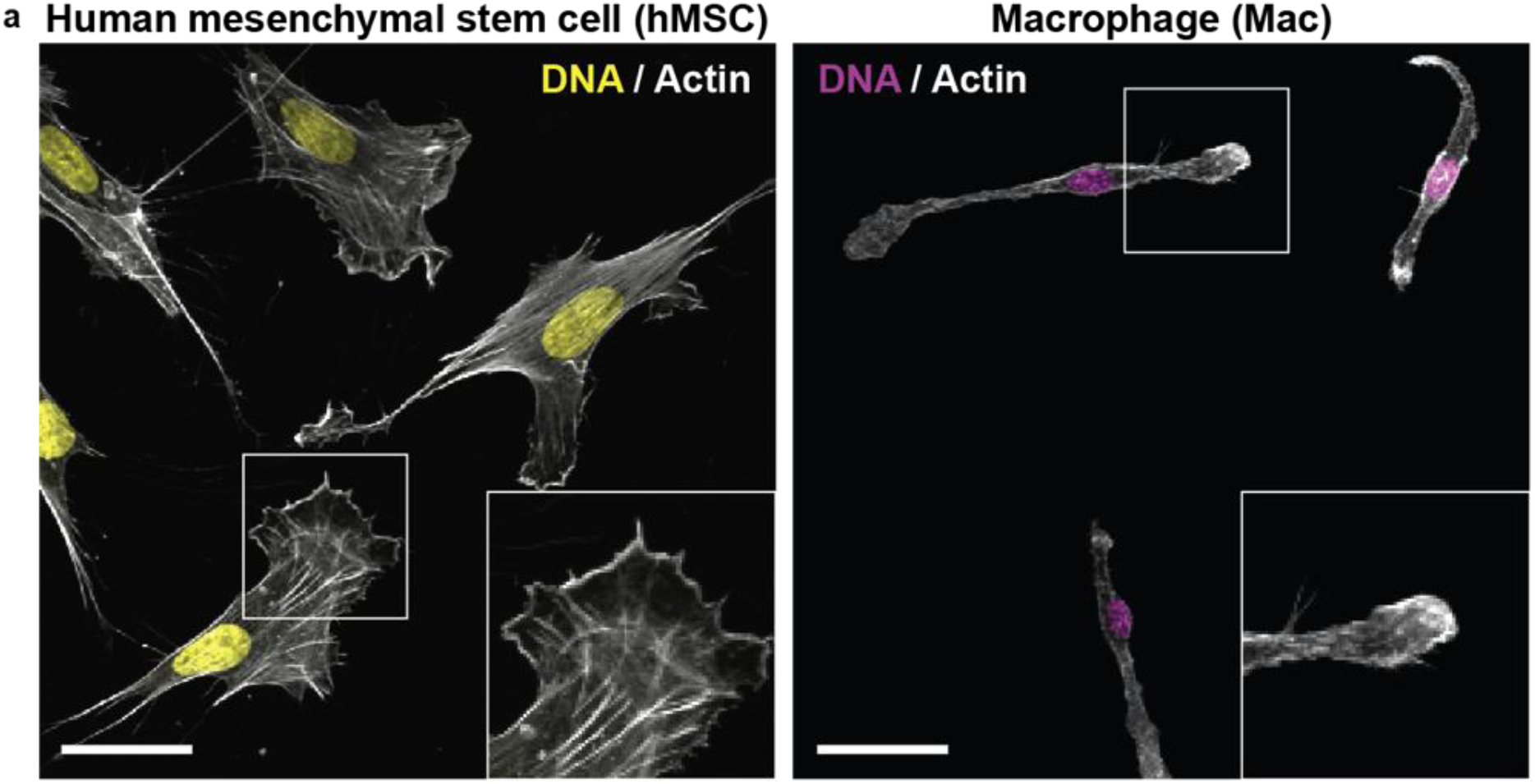
| Distinct cytoskeletal organization in hMSCs and macrophages. **a**, Representative images of hMSCs (left) and macrophages (right) morphology on PDMS. Insets highlight characteristic actin architectures, with hMSCs showing prominent stress fibers and broad lamellipodia, while macrophages display elongated, polarized morphologies with distinct actin-rich protrusions. Scale bars: 30 μm.

**Extended Data Fig. 7.**
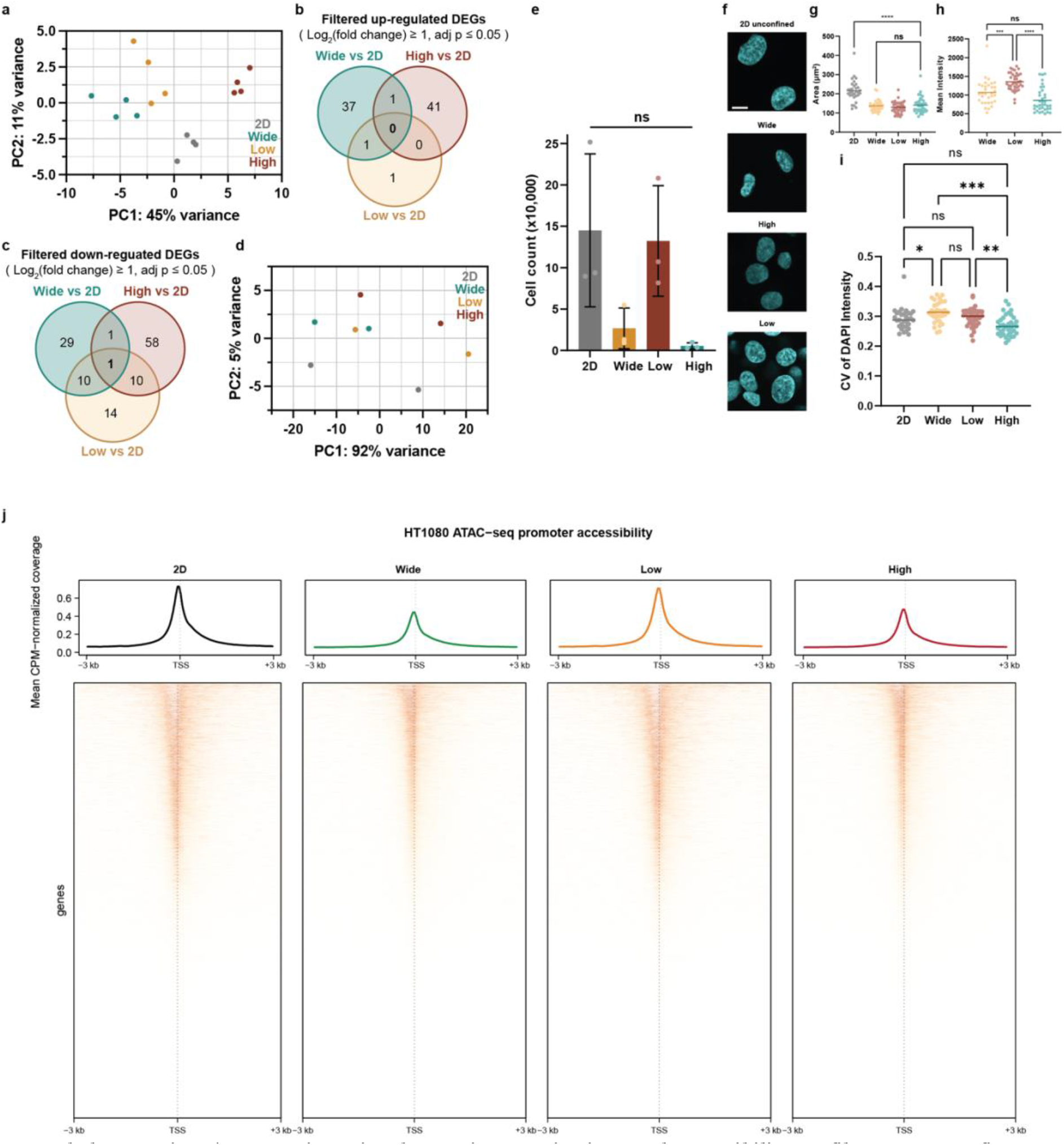
| Transcriptomic, chromatin organization, and accessibility profiles across confinement conditions. **a**, Principal component analysis (PCA) of RNA-seq data. **b**-**c**, Venn diagrams of upregulated (**b**) and downregulated (**c**) DEGs between confined and 2D conditions. **d**, PCA of ATAC-seq data. **e**, Quantification of cell yield under 2D, wide, low, and high confinement conditions. **f**, Representative maximum intensity projections of DAPI-stained nuclei of post-confinement HT1080 cells under 2D, wide, low-confinement, and high-confinement conditions. **g-i**, Quantification of nuclear area (**g**), mean nuclear DAPI intensity (**h**), and DAPI intensity heterogeneity using the coefficient of variation (CV, standard deviation/mean intensity) (**i**). Each dot represents one nucleus. ns, not significant; *p < 0.05, **p < 0.01, ***p < 0.001. Scale bars: 10 µm. **j**, ATAC-seq metaplots and heatmaps showing normalized chromatin accessibility centered on transcription start sites (TSS ± 3kb) under 2D, wide, low, and high confinement conditions.

**Extended Data Fig. 8.**
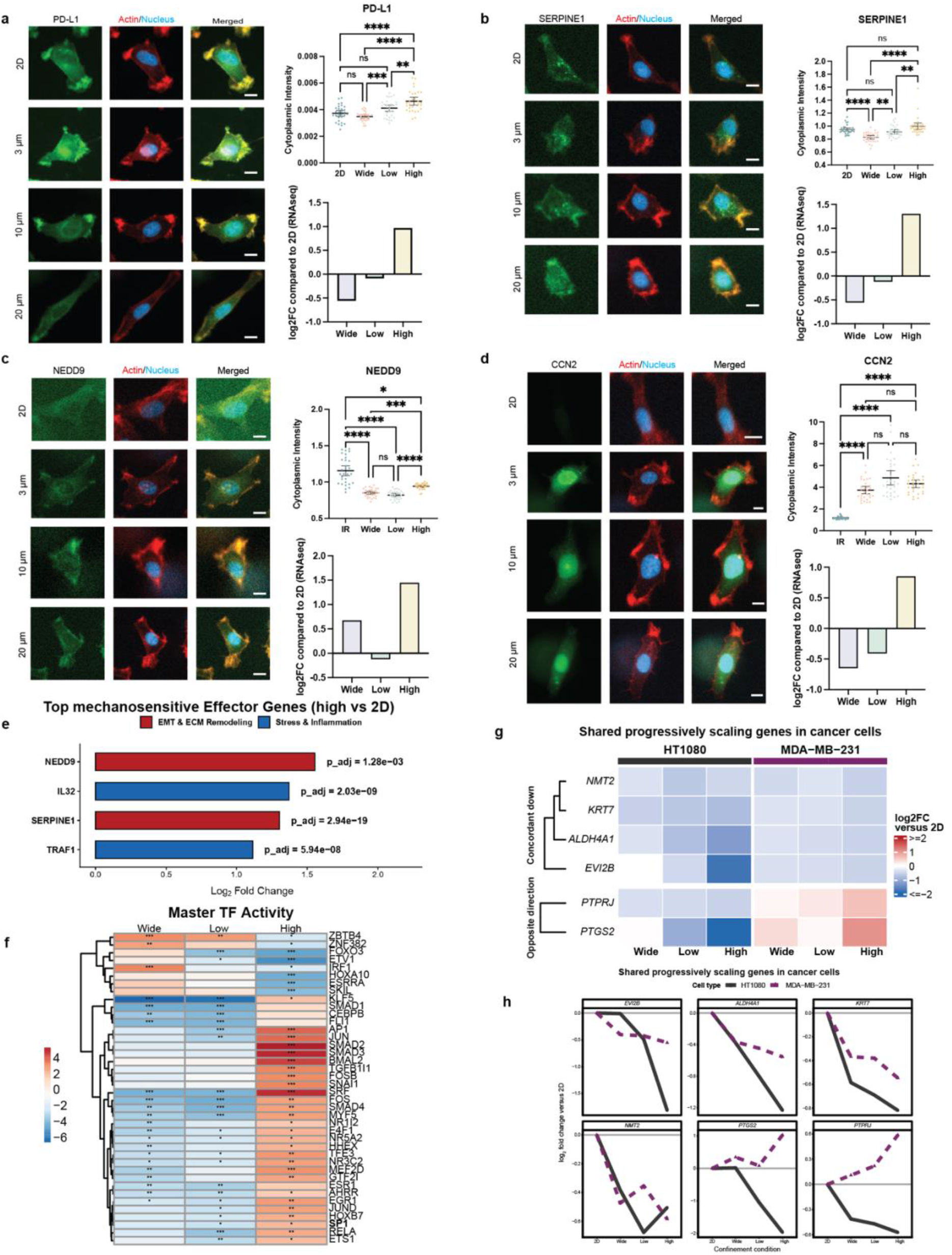
**| Validation and cross-cancer comparison of confinement-responsive transcriptional programs. a-d**, Representative immunofluorescence images and quantification of confinement-responsive targets PD-L1 (**a**), SERPINE1 (**b**), NEDD9 (**c**), and CCN2 (**d**) in cells under 2D, wide, low, and high confinement conditions. The confinement conditions were compared against 2D unconfined inner reservoir (IR) of the microchannel chip. (IR: n =30, Wide: n=30, Low: n=30, High: n=30). Statistical comparisons: one-way ANOVA with Tukey’s post-hoc test (a) and Kruskal-Wallis test with Dunn’s post-hoc test (b-d). Significance: ns (not significant), *P ≤ 0.05, **P ≤ 0.01, ***P ≤ 0.001, ****P ≤ 0.0001. Error bars: mean ± 95% C.I. Cytoplasmic fluorescence intensity is quantified alongside the corresponding RNA-seq log2 fold change relative to 2D controls. Scale bars: 10 µm. **e**, Top mechanosensitive effector genes identified in high confinement relative to 2D controls. **f**, Heatmap of inferred master transcription factor (TF) activity across wide, low, and high confinement conditions relative to 2D controls. *P ≤ 0.05, **P ≤ 0.01, ***P ≤ 0.001. **g**, Heatmap of genes displaying shared progressive confinement-dependent expression across HT1080 and MDA-MB-231 cancer cells. **h**, Expression profiles of the shared progressively scaling genes in HT1080 and MDA-MB-231 cells, showing changes in log2 fold change relative to 2D controls across increasing confinement.

**Extended Data Fig. 9.**
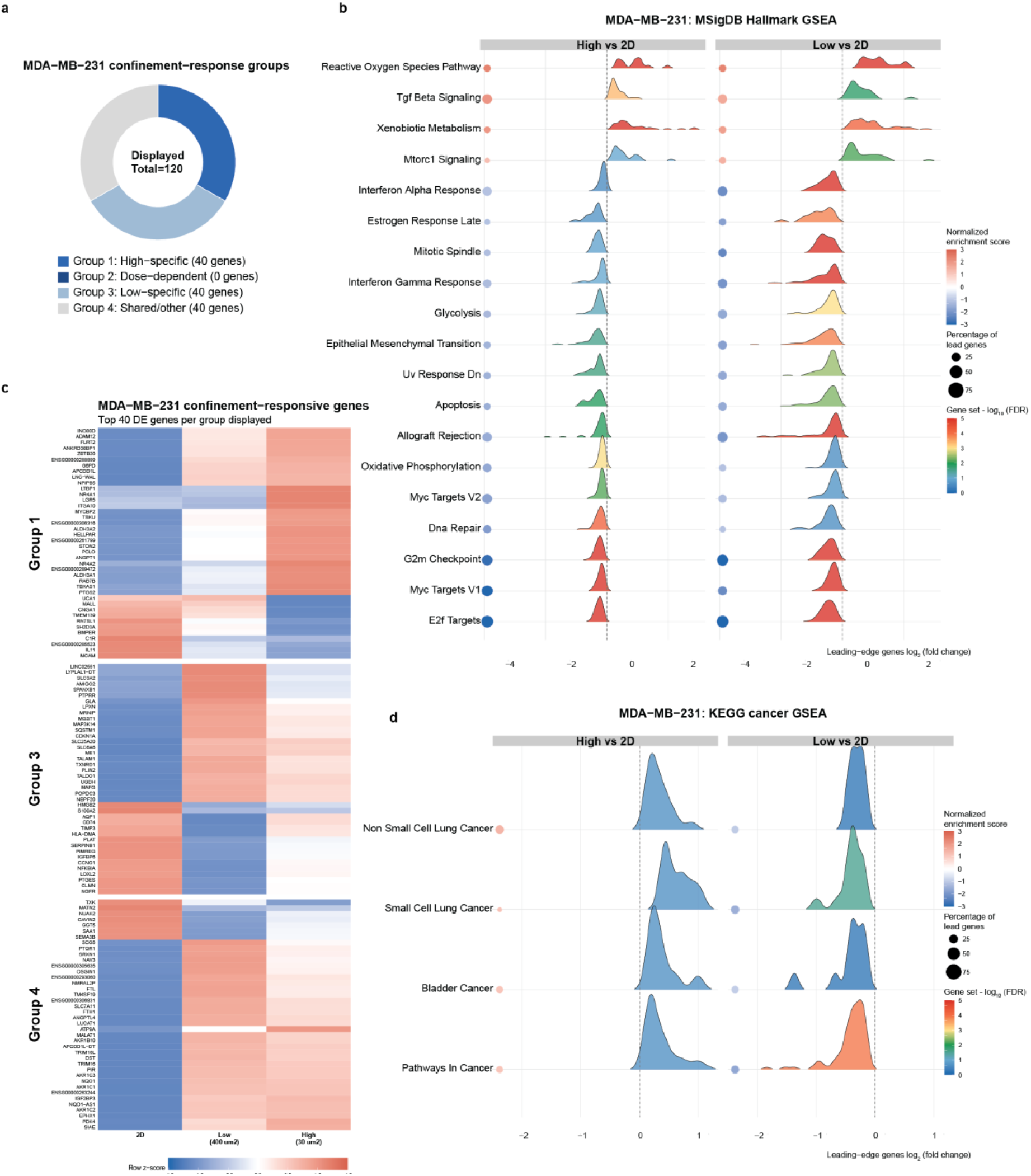
| Confinement-responsive transcriptomic programs in MDA-MB-231 cells. **a**, DEGs from high confinement vs 2D comparison in MDA-MB-231 cells were classified into four groups based on expression trends across 2D, low-confinement, and high-confinement conditions. **b**, GSEA analysis of MSigDB hallmark pathways. Enriched pathways are shown for high confinement vs 2D and low confinement vs 2D. **c**, Heatmap showing z-scored gene expression for Group 1, Group 3, and Group 4. No genes were classified into Group 2. **d**, GSEA analysis of KEGG cancer-related pathways. Enriched pathways are shown for high confinement vs 2D and low confinement vs 2D.

